# Hypothalamic Orexin Input to the Medial Amygdala Links Vigilance to Arousal

**DOI:** 10.64898/2026.03.09.710682

**Authors:** Xuaner Xiang, Chao Chen, Wei Zhou

**Author notes:** Corresponding author: Wei Zhou.

## Abstract

Arousal circuits govern anesthetic state transitions, but emergence is often complicated by agitation, and the neural circuits linking anesthetic arousal to vigilance remain unclear. Here, we identify a lateral hypothalamic orexin-to-medial amygdala pathway (LHA^Ox^→MeA) that links anesthetic state transitions to vigilance-like behavioral bias. Synaptic labeling and slice recordings revealed dense LHA^Ox^ innervation in the MeA, and RNAscope showed predominant Ox2R expression in MeA GABAergic (MeA^vGAT^) neurons. LHA^Ox^→MeA terminal activation increased local orexin release, promoted arousal from isoflurane anesthesia, delayed loss of righting reflex during induction, and accelerated recovery during emergence. In vigilance assays, stimulation acutely suppressed open-field exploration and produced real-time place preference without conditioned reinforcement. Fiber photometry demonstrated preferential recruitment of MeA^vGAT^ neurons across anesthetic states. Direct MeA^vGAT^ activation recapitulated cortical engagement, arousal, and vigilance phenotypes, whereas TeLC-mediated silencing of MeA^vGAT^ output abolished induction delay and reversed vigilance-like bias while sparing light-anesthesia arousal and emergence acceleration.

## Introduction

General anesthesia is a pharmacologically induced reversible state of unconsciousness that spans a continuum from light sedation to deep coma (1–4). Transitions into (induction) and out of anesthesia (emergence) represent critical brain-state changes that govern not only the loss and restoration of consciousness, but also how the brain re-engages with the environment (2, 5, 6). Clinically, emergence can often be complicated by agitation and dysregulated responsiveness to sensory input, underscoring that recovery of consciousness is not always behaviorally neutral. Despite recent advances in understanding arousal control, the circuits governing anesthesia states remain incompletely understood.

Arousal is a stimulus-evoked, phasic increase in alertness, whereas vigilance reflects the capacity to sustain alertness above a threshold and remain sensitive to environmental changes over time (7, 8). Vigilance-related monitoring can bias neural circuits toward rapid detection of salient or potentially threatening stimuli and lower the threshold for state transitions, and it can persist during reduced consciousness, including light sleep and anesthesia, and may oppose sleep entry and prime the brain for re-entry into wakefulness (5, 6, 9, 10). These considerations motivate the hypothesis that anesthetic arousal is coupled to limbic circuits that bias re-engagement toward vigilance-like monitoring, rather than uniformly restoring exploratory wake behavior. However, the circuit mechanisms that link anesthesia arousal to vigilance-like behavior during transitions into and out of unconsciousness remain poorly defined.

Orexin (Ox; hypocretin) neurons in the lateral hypothalamic area (LHA) are central regulators of arousal that integrate metabolic, emotional, and environmental cues to stabilize wakefulness and amplify motivated and stress-related behaviors (11, 12). They release orexin-A and orexin-B, and signal through the G-protein-coupled receptors Ox1R and Ox2R (13, 14). Loss of LHA^Ox^ signaling underlies narcolepsy with cataplexy (15–18). Ox neurons project widely to coordinate diverse functions, including basal forebrain and locus coeruleus supporting wakefulness and cortical activation (19, 20), paraventricular thalamus and ventral tegmental area modulating motivation and reward-related behaviors (21–23), and medullary targets regulating autonomic function (24–26). This broad architecture suggests that orexin’s arousal drive may be routed through discrete downstream nodes. However, which limbic nodes couple orexin-driven arousal to vigilance-related behavioral bias remains unclear.

We identified prominent orexin projections to the amygdala, with particularly dense innervation of the medial amygdala (MeA) (27, 28). The amygdala coordinates adaptive behavioral and physiological responses to emotionally salient stimuli (29, 30) and contributes to vigilance by evaluating environmental uncertainty, internal state, and stimulus salience (31, 32). MeA is uniquely positioned to integrate internal state signals with salience detection, given its strong hypothalamic and olfactory-related inputs and its outputs to autonomic and neuroendocrine centers (33). Moreover, reduced responsiveness during sleep and anesthesia does not imply uniform silencing of brain circuits (34). The MeA may remain positioned to couple arousal drive to vigilance-related processing across anesthetic state transitions.

Here, we defined the LHA^Ox^→MeA projection that links arousal control with vigilance behavior bias. Using viral tracing, electrophysiology, and RNAscope in situ hybridization, we showed that LHA^Ox^ neurons form functional synaptic connections in the MeA and identify MeA vGAT^+^ neurons (MeA^vGAT^) as a key downstream substrate. Combining optogenetics with EEG/EMG and fiber photometry, we tested how LHA^Ox^→MeA signaling regulates anesthetic state transitions and biases vigilance-related behavior. Finally, we showed that MeA^vGAT^ activation is sufficient to recruit prefrontal cortical activity and recapitulate arousal and vigilance-like phenotypes, whereas silencing MeA^vGAT^ synaptic output selectively abolishes orexin-dependent induction delay and disrupts vigilance bias, establishing MeA^vGAT^ output as a key relay that translates orexin input into cortical and behavioral consequences.

## Results

### LHA^Ox^ neurons form functional synaptic connections in the MeA

To determine whether LHA^Ox^ neurons innervate the medial amygdala (MeA), we used a synapse-labeling strategy in Orexin-Cre mice by injecting Cre-dependent AAV1-FLEX-tdTomato-T2A-SypEGFP into the LHA to label orexin axons and presynaptic terminals. We found that synaptophysin-EGFP-positive puncta were abundant within the MeA, indicating dense terminal labeling consistent with synapse-like contacts from LHA^Ox^ neurons (Fig. 1A–C).

**Fig. 1.**
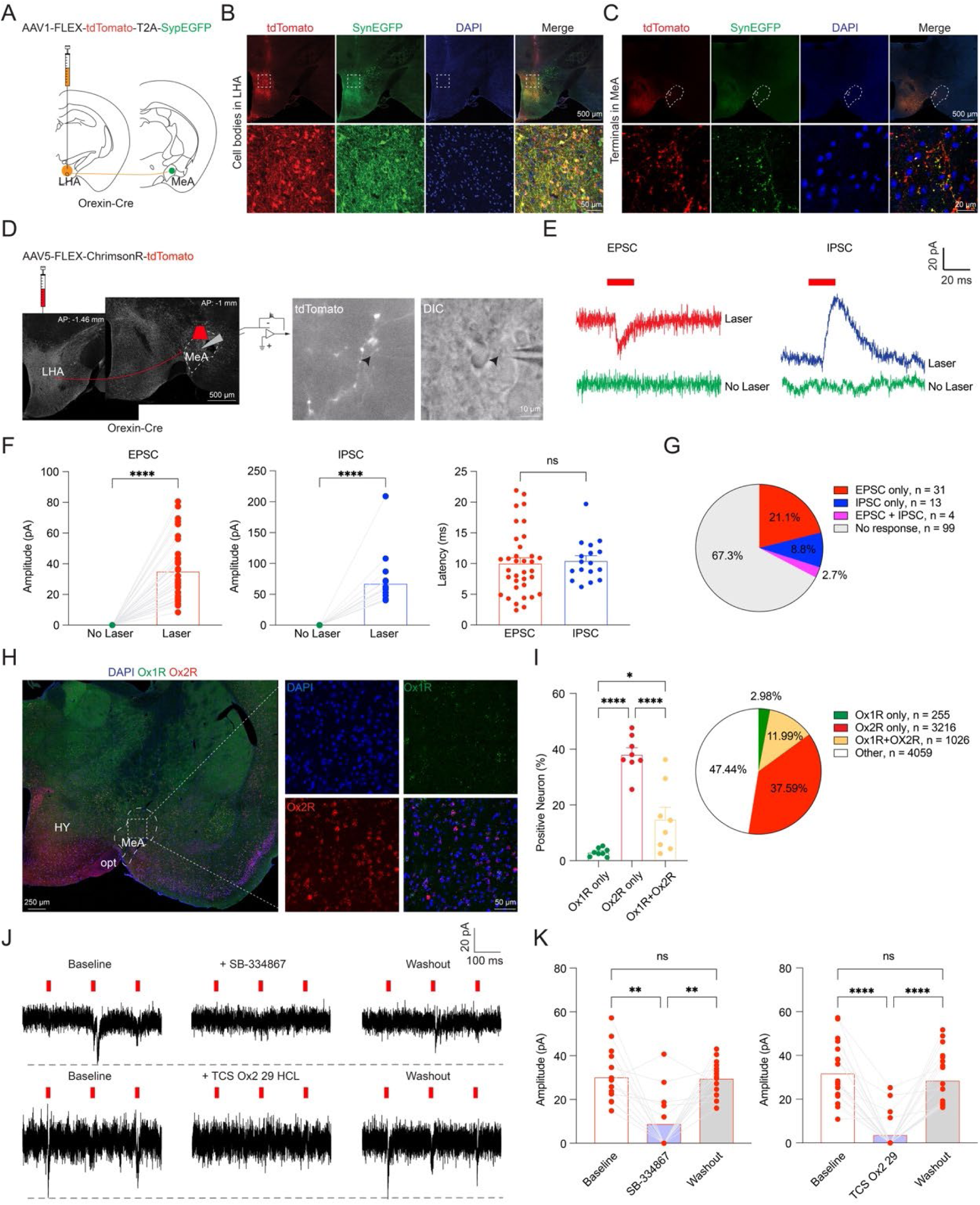
LHA^Ox^ neurons form functional synaptic connections in the medial amygdala. (**A**) Cre-dependent AAV1-FLEX-tdTomato-T2A-SynpEGFP was injected into the lateral hypothalamic area (LHA) of Orexin-Cre mice to label orexin terminals in the medial amygdala (MeA). (**B** to **C**) Representative images showing orexin neuron somata in the LHA (**B**) and dense LHA^Ox^ synaptic terminal labeling in the MeA (**C**). (**D**) Cre-dependent AAV5-FLEX-ChrimsonR-tdTomato was injected into the LHA of Orexin-Cre mice, and whole-cell recordings were obtained from MeA neurons (left); tdTomato^+^ orexin fibers surround the recorded neuron (right). (**E** to **F**) Optical stimulation (635 nm) of LHA^Ox^ terminals evoked both excitatory (EPSCs) and inhibitory (IPSCs) postsynaptic currents in MeA neurons; group quantification is shown. (**G**) Distribution of electrophysiologically defined MeA neuron subtypes recorded. (**H** to **I**) RNAscope detection of Ox1R and Ox2R mRNA in the MeA (**H**) with quantification of receptor-expressing populations (**I**); pie chart summarizes pooled cell-type proportions. (**J** to **K**) Pharmacological blockade of orexin receptors suppressed LHA^Ox^ terminal–evoked EPSCs: responses were reduced by the Ox1R antagonist SB-334867 or the Ox2R antagonist TCS OX2 29 and recovered after washout; quantification is shown. Scale bars as indicated. Data are mean ± S.E.M.; statistical details and exact n values are provided in Supplementary Table 1. *p ≤ 0.05, **p ≤ 0.01, ****p < 0.0001; LHA, lateral hypothalamic area; MeA, medial amygdalar nucleus; HY, hypothalamus; opt, optic tract.

We next assessed whether these projections are functionally active. We expressed the red-shifted opsin ChrimsonR in LHA^Ox^ neurons (AAV5-FLEX-ChrimsonR-tdTomato) and performed whole-cell recordings from MeA neurons in acute brain slices (Fig. 1D). TdTomato-positive orexin fibers surrounded recorded MeA neurons, and optical stimulation of LHA^Ox^ terminals reliably evoked both excitatory and inhibitory postsynaptic currents (EPSCs and IPSCs), indicating that LHA^Ox^ terminal activation engages local MeA microcircuits (Fig. 1E). EPSC and IPSC onset latencies were in a similar range, consistent with rapid synaptic transmission within the recruited circuit (Fig. 1F). Recordings spanned multiple electrophysiologically defined MeA neuron subtypes, suggesting broad targeting across heterogeneous neuronal populations (Fig. 1G).

To define the receptor landscape supporting orexin signaling in the MeA, we performed RNAscope for Ox1R and Ox2R mRNAs (Fig. 1H). Ox2R expression predominated over Ox1R, with a subset of cells co-expressing both receptors (Fig. 1H–I), indicating that Ox2R is the major orexin receptor in this region.

Consistent with this distribution, pharmacological blockade of orexin receptors attenuated terminal stimulation–evoked EPSCs in MeA neurons (Fig. 1J–K). The Ox1R antagonist SB-334867 reduced EPSC amplitude, whereas the Ox2R antagonist TCS Ox2 29 produced a more pronounced suppression. EPSCs recovered toward baseline after washout, supporting receptor-specific and reversible pharmacological effects. In contrast, IPSCs were not significantly affected by either antagonist (Supplementary Fig. 1B), suggesting that the inhibitory responses evoked by terminal stimulation reflect recruitment of local inhibitory circuitry rather than direct orexin receptor–mediated inhibition.

Together, these data demonstrate that LHA^Ox^ neurons form a dense, functional projection to the MeA, engage multiple local neuronal subtypes, and signal predominantly through Ox2R.

### The LHA^Ox^→MeA projection selectively regulates arousal and anesthetic transitions

To investigate whether the LHA^Ox^→MeA projection participates in anesthetic state transitions, we selectively manipulate MeA-projecting LHA^Ox^ neurons in vivo by injecting Cre-dependent retrograde AAVrg-DIO-ChR2-mCherry into the MeA of Orexin-Cre mice and implanting optic fibers together with EEG/EMG electrodes (Fig. 2A–B). Optogenetic activation increased cFos expression in MeA neurons relative to mCherry controls, confirming engagement of local MeA circuitry by LHA^Ox^ inputs (Fig. 2C–D).

**Fig. 2.**
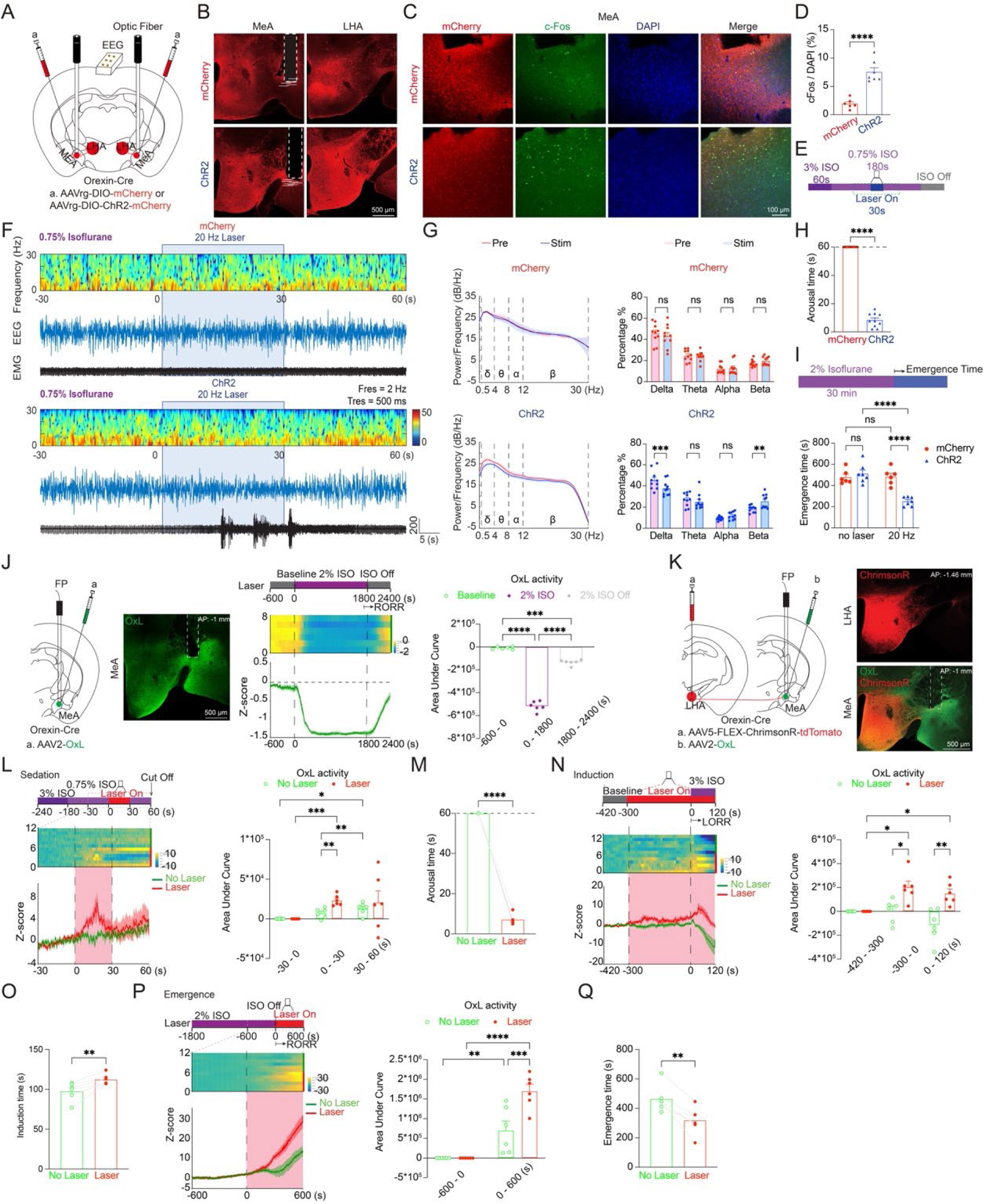
Activation of the LHA^Ox^→MeA circuit promotes arousal and modulates anesthetic state transitions via orexin release. (**A**) Retrograde targeting strategy: Cre-dependent AAVrg-DIO-ChR2-mCherry was injected into the MeA of Orexin-Cre mice to label LHA^Ox^ neurons projecting to the MeA, followed by implantation of optic fibers and EEG/EMG electrodes. (**B**) Representative images confirming viral expression in MeA and LHA; horizontal bars indicate individual fiber tip locations. Scale bars, 500 µm. (**C** to **D**) Optogenetic activation increased neuronal activity in the MeA, assessed by cFos immunostaining and quantification (**C**, representative; **D**, summary). (**E**) Schematic of the isoflurane sedation arousal paradigm. (**F** to **H**) Under 0.75% isoflurane, 20 Hz stimulation of the LHA^Ox^→MeA pathway increased cortical activation and promoted behavioral arousal in ChR2 mice but not mCherry controls (**F**, representative EEG/EMG and spectrogram; **G**, EEG power spectral density and band power analysis; **H**, arousal latency). (**I**) Schematic of the emergence paradigm and quantification of righting recovery: LHA^Ox^→MeA activation accelerated recovery of the righting reflex after deep anesthesia in ChR2 mice but not in controls. (**J**) Fiber photometry of orexin release in the MeA using the genetically encoded orexin sensor OxLight (OxL): OxL signal decreased during 2% isoflurane anesthesia and increased during emergence; AUC analysis summarizes state-dependent changes. (**K**) Strategy for simultaneous LHA^Ox^ terminal stimulation and MeA orexin sensing (ChrimsonR in LHA; OxL in MeA) with representative expression images. (**L** to **M**) During isoflurane sedation, LHA^Ox^→MeA terminal stimulation increased MeA orexin release (**L**) and induced rapid arousal from light anesthesia (**M**). (**N** to **O**) During induction, pre-activation of LHA^Ox^→MeA terminals increased MeA orexin release (**N**) and delayed loss of the righting reflex (**O**). (**P** to **Q**) During emergence, terminal stimulation accelerated MeA orexin release (**P**) and shortened the time to recovery of the righting reflex (**Q**). Data are mean ± S.E.M.; statistical details and exact n values are provided in Supplementary Table 1. *p ≤ 0.05, **p ≤ 0.01, ****p < 0.0001.

Sleep and general anesthesia share overlapping arousal-regulating circuits, and orexin is widely recognized as a key modulator of sleep-wake stability and arousal (6). We first tested whether activating the LHA^Ox^→MeA projection can promote awakening from non-rapid eye movement (NREM) sleep. We found that during stable NREM sleep, 20 Hz stimulation induced robust EEG-defined state transitions in ChR2 mice but not in controls (Supplementary Fig. 2A). Spectral analysis was consistent with cortical activation, with reduced delta power and increased higher-frequency activity, and stimulation increased behavioral arousal from NREM sleep across frequencies (Supplementary Fig. 2B–C). It was consistent with our prior work that revealed the orexin projections to the substantia innominata mediate arousal in NREM sleep and anesthesia (35). In contrast to prior findings that global activation of LHA^Ox^ neurons can produce analgesia (28, 36–39), selective LHA^Ox^→MeA activation did not change hot-plate withdrawal latency, suggesting that this projection is not sufficient to modulate thermal nociception under these conditions (Supplementary Fig. 2D).

Building on our prior work establishing a light isoflurane protocol that maintains mice in a sedated state and enables clear observation of orexin-driven arousal behaviors (28). We next tested whether LHA^Ox^→MeA activation promotes arousal from anesthetic sedation and accelerates emergence. Under 0.75% isoflurane sedation, 20 Hz stimulation reliably induced behavioral arousal in ChR2 mice (mean latency ∼8 s) and produced EEG changes consistent with cortical activation, whereas mCherry controls showed no comparable effect (Fig. 2F–H). During emergence from deep anesthesia (2% isoflurane), stimulation significantly shortened the latency to recovery of the righting reflex (RoRR) in ChR2 mice but not in controls (Fig. 2I). Thus, selective activation of MeA-projecting LHA^Ox^ neurons promotes arousal from sedation and facilitates emergence.

A prior study reported that orexin depolarized amygdala neurons and increased their firing via orexin receptors (40). We therefore hypothesized that LHA^Ox^→MeA activation promotes arousal from anesthetic sedation and accelerates emergence by increasing orexin release in the MeA to excite local MeA neurons. To directly test this, we expressed the genetically encoded orexin sensor OxLight (OxL) in the MeA and performed fiber photometry to track MeA orexin dynamics across anesthetic transitions. We focused on induction, steady-state sedation, and emergence because these phases represent distinct operational “checkpoints” of anesthetic state control. Induction marks entry into unconsciousness, sedation reflects a relatively stable anesthetic state, and emergence marks recovery from unconsciousness. These transitions may not be controlled by the same neural mechanisms, and orexin signaling has been particularly implicated in emergence (6, 41, 42).

OxL fluorescence declined during 2% isoflurane anesthesia and increased during spontaneous emergence, indicating state-dependent modulation of orexin release in the MeA (Fig. 2J). To test causal control, we combined MeA OxL expression with optogenetic excitation (ChrimsonR, Fig. 2K) or inhibition (Jaws, Supplementary Fig. 3A) of LHA^Ox^ neurons. Under light anesthesia, terminal activation increased MeA OxL signal and induced rapid arousal (Fig. 2L–M). During induction, pre-activation elevated MeA OxL signal and delayed loss of the righting reflex (LoRR) (Fig. 2N–O). During emergence, stimulation increased MeA OxL signal and accelerated RoRR (Fig. 2P–Q).

In contrast, optogenetic inhibition of the LHA^Ox^→MeA projection did not further reduce MeA OxL signal in the awake baseline and did not significantly alter arousal from sedation, LoRR during induction, or RoRR during emergence (Supplementary Fig. 3B–G). Although inhibition decreased MeA OxL signal during induction epochs, this reduction was not sufficient to shorten induction latency (Supplementary Fig. 3D–E). The efficacy of Jaws-mediated suppression was confirmed ex vivo by whole-cell recordings from Jaws-expressing LHA^Ox^ neurons, in which optical stimulation hyperpolarized membrane potential and blocked evoked spiking (Supplementary Fig. 3H–I).

Together, these results show that orexin release in the MeA is dynamically regulated across induction, sedation, and emergence, and that activation of the LHA^Ox^→MeA projection is sufficient to elevate local orexin release and modulate transitions into and out of anesthesia.

### LHA^Ox^ → MeA projection activation biases vigilance-like behavior and real-time preference without inducing conditioned reinforcement

Given that vigilance encompasses arousal, and that both orexin signaling and amygdala circuits promote high-alert, threat-responsive behavioral states (43–46), we asked whether LHAOx→MeA activation drives anesthesia arousal and biases vigilance-like behavior. Mice expressing mCherry or ChrimsonR in the LHA orexin neurons with the optic fibers implanted above the MeA (Fig.3A) were subjected to the open field test (OFT, Fig.3B) to quantify exploration and vigilance-like avoidance, real-time place preference (RTPP, Fig.3E) to measure the immediate valence of LHA^Ox^→MeA projection activation, and conditioned place preference (CPP, Fig.3H) to test whether repeated activation produces a lasting learned preference.

**Fig. 3.**
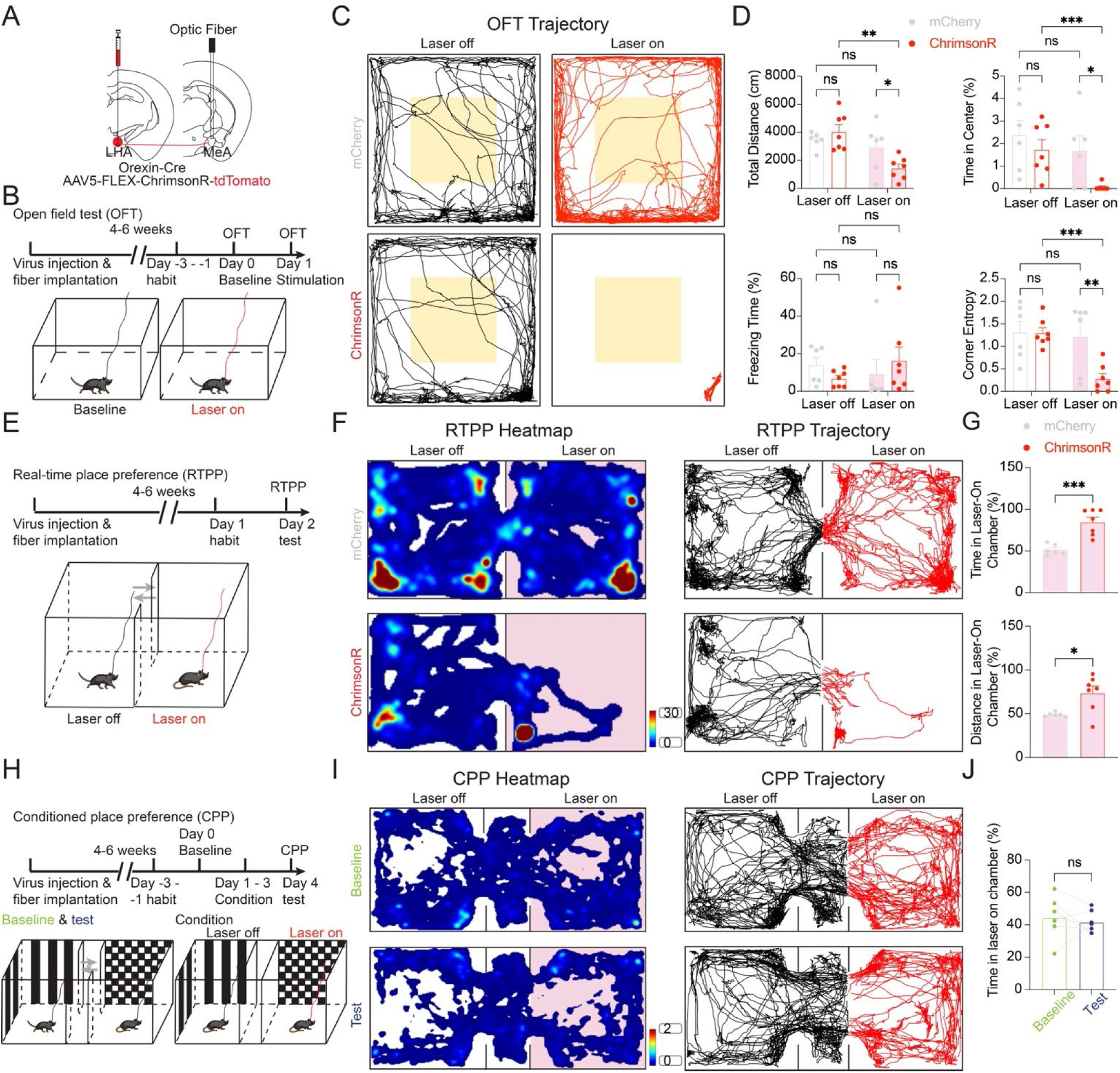
Activation of the LHA^Ox^→MeA projection acutely suppresses exploration and biases real-time preference without conditioned reinforcement. (**A**) Cre-dependent AAV5-FLEX-ChrimsonR-tdTomato was injected into the LHA of Orexin-Cre mice, and an optic fiber was implanted above the MeA. (**B**) Experimental timeline and schematic of the open field test (OFT). (**C** to **D**) Representative OFT trajectories (**C**: mCherry, gray; ChrimsonR, red; laser-off and laser-on epochs; center zone outlined) and quantification of total distance traveled, percent time in the center, freezing time, and corner entropy (**D**). (**E**) Experimental timeline and schematic of real-time place preference (RTPP). (**F** to **G**) Representative RTPP heatmaps/trajectories (**F**) and quantification showing that LHA^Ox^→MeA terminal stimulation increased time spent and distance traveled in the laser-paired chamber in the ChrimsonR group but not in mCherry controls (**G**). (**H**) Experimental timeline and schematic of conditioned place preference (CPP). (**I** to **J**) Representative CPP heatmaps/trajectories from the same animal during baseline and test sessions (**I**) and quantification showing no change in time spent in the stimulation-paired chamber following conditioning (**J**). Data are mean ± S.E.M.; statistical details and exact n values are provided in Supplementary Table 1. *p ≤ 0.05, **p ≤ 0.01, ***p < 0.001.

In the OFT, two-way ANOVA (group x laser) revealed a significant interaction for total distance traveled (Fig.3D), and post hoc comparisons showed that ChrimsonR mice exhibited reduced locomotion during laser-on conditions compared with laser-off conditions and controls. For the percentage of the time in the center, planned comparisons indicated a reduction in ChrimsonR mice during laser-on conditions, but not laser-off conditions. Corner entropy was lower in ChrimsonR mice during laser-on than laser-off conditions, and this reduction was also significant relative to control mice, indicating more stereotyped and restricted corner occupancy. These effects were observed within ChrimsonR mice comparing laser-off versus laser-on, whereas the mCherry group showed no significant stimulation-dependent differences. Notably, total freezing time was not increased, indicating that the reduced distance or center exploration was not attributable to immobility.

We next asked whether this behavioral bias was accompanied by an immediate preference for the stimulated context. In the RTPP assay, activation of the LHA^Ox^→MeA projection produced a robust real-time preference in the ChrimsonR group, as evidenced by increased time spent in the laser-on chamber (Fig.3G) and the distance traveled in the laser-on chamber (Fig.3G). Notably, the increased distance was largely driven by repetitive corner-confined back-and-forth turning rather than broad exploration across the chamber, suggesting a vigilance-like locomotor pattern in the laser-on chamber.

In contrast, when we paired repeated activation of the LHA^Ox^→MeA projection with a specific chamber during conditioning, this manipulation failed to induce a lasting place preference. CPP testing revealed no difference in time spent in the stimulation-paired chamber between baseline and test sessions (Fig.3J).

Together, these results suggested that activation of the LHA^Ox^→MeA projection acutely suppressed the exploration and promoted vigilance-like behavior while biasing real-time place preference, but is insufficient to produce a lasting conditioned effect.

### LHA^Ox^ inputs preferentially engage MeA^vGAT^ to regulate anesthetic arousal and state transitions

To identify candidate downstream targets through which the LHA^Ox^→MeA projection regulates anesthetic arousal, we mapped Ox1R and Ox2R expression in the MeA using RNAscope together with markers for major neurotransmitter classes (vGAT, vGlut2, and TH), guided by the Allen Institute cell-type atlas. Ox2R expression predominated in the MeA, and vGAT^+^ inhibitory neurons comprised the largest neuronal population in this region (Fig. 4A–B; Supplementary Fig. 4A–D). Co-expression analysis further indicated that both inhibitory and excitatory MeA populations contain orexin receptor–positive subsets, with Ox2R present in a larger fraction of receptor-positive cells across vGAT^+^ and vGlut2^+^ classes (Fig. 4B; Supplementary Fig. 4B, D).

**Fig. 4.**
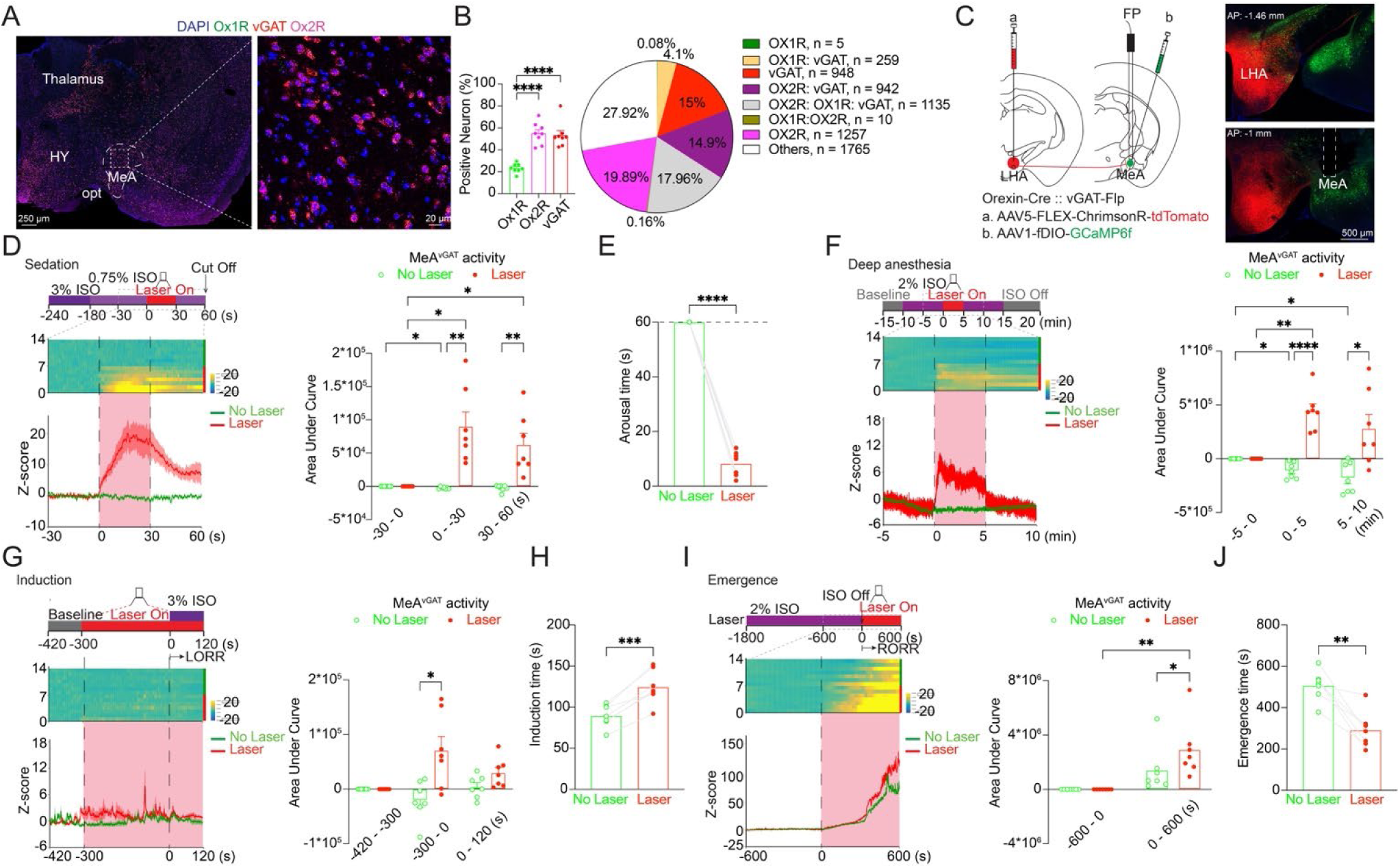
LHA^Ox^→MeA activation recruits MeA^vGAT^ neurons to promote arousal and modulate anesthetic state transitions. (**A** to **B**) RNAscope detection of Ox1R and Ox2R mRNA in MeA neurons together with vGAT, with quantification of receptor- and vGAT-expressing populations (**A**, representative; **B**, summary with pooled distribution). (**C**) Intersectional strategy to stimulate LHA^Ox^ terminals (Cre-dependent ChrimsonR in LHA^Ox^) while recording MeA^vGAT^ activity (Flp-dependent GCaMP6f in MeA^vGAT^) in Orexin-Cre::vGAT-Flp mice; representative images confirm expression and fiber placement. (**D** to **F**) Fiber photometry showing that LHA^Ox^ terminal activation increases MeA^vGAT^ activity during isoflurane sedation (**D**) and during 2% isoflurane anesthesia (**F**), and induces rapid arousal from light anesthesia (**E**). (**G** to **H**) During induction, pre-activation of LHA^Ox^ terminals elevates MeA^vGAT^ activity (**G**, including AUC analysis) and delays loss of the righting reflex (LoRR) (**H**). (**I** to **J**) During emergence, LHA^Ox^ terminal activation increases MeA^vGAT^ activity (**I**) and accelerates recovery of the righting reflex (RoRR) (**J**). Data are mean ± S.E.M.; statistical details and exact n values are provided in Supplementary Table 1. *p ≤ 0.05, **p ≤ 0.01, ****p < 0.0001.

We next tested which MeA neuronal class is recruited during orexin-driven anesthetic arousal using intersectional photometry. We expressed Cre-dependent ChrimsonR in LHA^Ox^ neurons and Flp-dependent GCaMP6f in either MeA^vGlut2^ neurons (Orexin-Cre::vGlut2-Flp) or MeA^vGAT^ neurons (Orexin-Cre::vGAT-Flp) (Fig. 4C; Supplementary Fig. 4E). Under isoflurane sedation, LHA^Ox^→MeA terminal stimulation reliably induced behavioral arousal but did not produce a detectable change in MeA^vGlut2^ population activity (Supplementary Fig. 4F–G). Consistent with this functional result, Flp-dependent labeling in the MeA of Orexin-Cre::vGlut2-Flp mice was sparse. Whereas the control injections of a Flp-dependent reporter/effector into vGlut2-enriched regions (diagonal band nucleus (NDB) and hypothalamus (HY)) produced robust expression, supporting efficient Flp-dependent expression in this line and indicating relatively low vGlut2+ representation in the MeA sampling region (Supplementary Fig. 4H). Together, these findings suggested that MeA^vGlut2^ neurons are not robustly recruited during LHA^Ox^→MeA stimulation in the context of anesthesia arousal.

In contrast, in Orexin-Cre::vGAT-Flp mice, LHA^Ox^→MeA terminal stimulation robustly increased MeA^vGAT^ activity during isoflurane sedation and induced rapid arousal from light anesthesia (Fig. 4D–E). MeA^vGAT^ activity also increased during deep anesthesia (2% isoflurane) (Fig. 4F). During induction, pre-activation elevated MeA^vGAT^ activity relative to time-matched no-laser controls during the pre-induction window, and this was accompanied by a significant delay in loss of the righting reflex (LoRR) (Fig. 4G–H). During emergence, stimulation increased MeA^vGAT^ activity and significantly accelerated recovery of the righting reflex (RoRR) (Fig. 4I–J). These results identify MeA^vGAT^ neurons as a key downstream population engaged by LHA^Ox^ input across anesthetic states.

To control for nonspecific effects of light delivery and photometry, we performed parallel experiments in mCherry controls. Optical stimulation did not alter MeA^vGAT^ activity across induction, sedation, deep anesthesia, or emergence, and did not affect arousal latency, LoRR, or RoRR (Supplementary Fig. 5A–H).

### MeA^vGAT^ neurons act as a state-dependent gate for the LHA^Ox^-driven arousal and behavioral valence

To determine whether MeA^vGAT^ activation is sufficient to promote anesthetic arousal, we expressed Flp-dependent ChrimsonR in the MeA of vGAT-Flp mice and monitored prelimbic cortex (PrL) activity with GCaMP6s (Fig. 5A). We monitored PrL because prefrontal cortical activation is a well-established marker of arousal state and provides a sensitive readout of anesthesia-to-wake transitions (47). During isoflurane sedation, MeA^vGAT^ activation increased PrL activity and reliably induced behavioral arousal (Fig. 5B–C). Under deep anesthesia (2% isoflurane), MeA^vGAT^ activation did not significantly elevate PrL activity (Fig. 5D), consistent with strong anesthetic suppression of cortical dynamics.

**Fig. 5.**
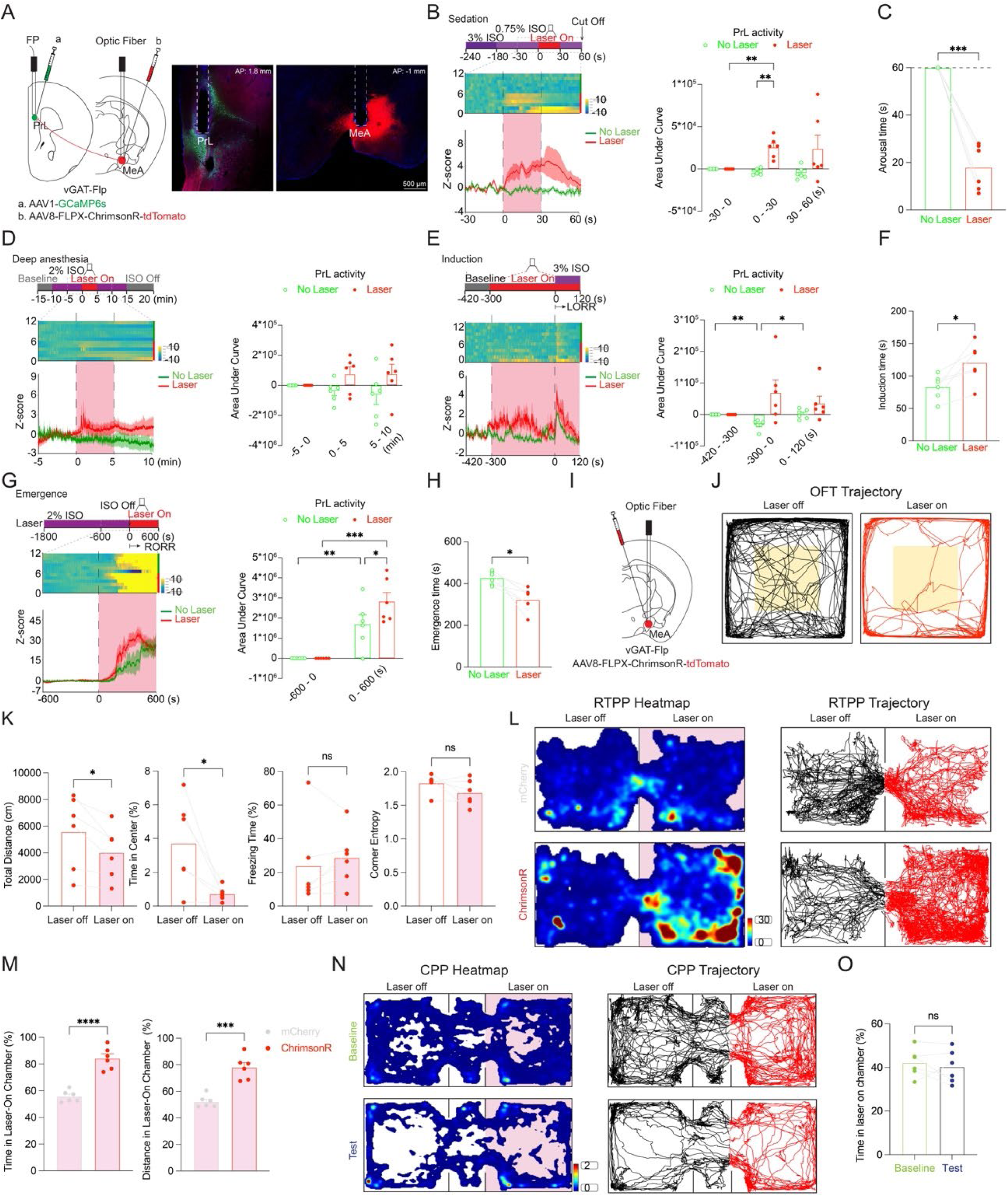
Activation of MeA^vGAT^ neurons drives cortical engagement, modulates anesthetic state transitions, and biases vigilance-like behavior. (**A**) Strategy to activate MeA^vGAT^ neurons (Flp-dependent ChrimsonR in MeA^vGAT^) while monitoring prelimbic cortex (PrL) activity (AAV1-GCaMP6s in PrL) in vGAT-Flp mice; representative images confirm expression and fiber placement. (**B**, **D**) Fiber photometry showing PrL activity during MeA^vGAT^ activation under isoflurane sedation (**B**) and 2% isoflurane anesthesia (**D**), with quantification. (**C**) MeA^vGAT^ activation induces behavioral arousal from light anesthesia. (**E** to **F**) During induction, pre-activation of MeA^vGAT^ neurons modulates PrL activity (**E**) and delays loss of the righting reflex (LoRR) (**F**). (**G** to **H**) During emergence, MeA^vGAT^ activation increases PrL activity (**G**) and accelerates recovery of the righting reflex (RoRR) (**H**). (**I**) Strategy for behavioral assays: viral expression and optic fiber implantation above the MeA in vGAT-Flp mice. (**J** to **K**) Representative open-field trajectories (**J**) and quantification of exploration and defensive metrics (**K**). (**L** to **M**) Representative RTPP heatmaps/trajectories (**L**) and quantification showing increased time spent and distance traveled in the laser-paired chamber in ChrimsonR mice but not mCherry controls (**M**). (**N** to **O**) Representative CPP heatmaps/trajectories during baseline and test sessions (**N**) and quantification showing no change in time spent in the stimulation-paired chamber following conditioning (**O**). Data are mean ± S.E.M.; statistical details and exact n values are provided in Supplementary Table 1. *p ≤ 0.05, **p ≤ 0.01, ***p < 0.001, ****p < 0.0001.

During induction, PrL activity showed time-dependent drift in no-laser controls, and stimulation did not produce a consistent increase in PrL activity across the pre-induction window (Fig. 5E). Nevertheless, MeA^vGAT^ pre-activation significantly delayed loss of the righting reflex (LoRR) (Fig. 5F). During emergence, MeA^vGAT^ activation increased PrL activity and significantly accelerated recovery of the righting reflex (RoRR) (Fig. 5G–H). Together, these results indicate that MeA^vGAT^ activation is sufficient to promote arousal and to bidirectionally modulate anesthetic state transitions by delaying induction while accelerating emergence.

We next asked whether MeA^vGAT^ activation is sufficient to recapitulate the vigilance-like behavioral bias observed with LHA^Ox^→MeA stimulation. In the OFT test, MeA^vGAT^ stimulation reduced locomotor exploration and center engagement without increasing freezing time (Fig. 5K), consistent with the effects of LHA^Ox^→MeA terminal activation in Fig. 3D. However, it did not produce the prominent corner-restricted locomotor pattern observed during LHA^Ox^→MeA terminal activation. In the RTPP test, MeA^vGAT^ stimulation increased both time spent and distance traveled in the laser-paired compartment (Fig. 5M). Notably, this preference was accompanied by perimeter-biased locomotion, consistent with a distributed thigmotactic “patrolling” mode rather than the corner-confined pattern observed during LHA^Ox^→MeA terminal stimulation. In contrast, MeA^vGAT^ stimulation did not produce conditioned reinforcement, as the CPP test showed no preference shift relative to baseline (Fig. 5O).

To test whether MeA^vGAT^ output is required for LHA^Ox^→MeA-driven cortical engagement and state modulation, we blocked synaptic output from MeA^vGAT^ neurons with Flp-dependent TeLC while expressing Cre-dependent ChrimsonR in LHA^Ox^ neurons and monitoring PrL activity (Fig. 6A). TeLC blocks synaptic transmission by cleaving synaptobrevin/VAMP2 to prevent vesicular neurotransmitter release from infected neurons (48–50).

**Fig. 6.**
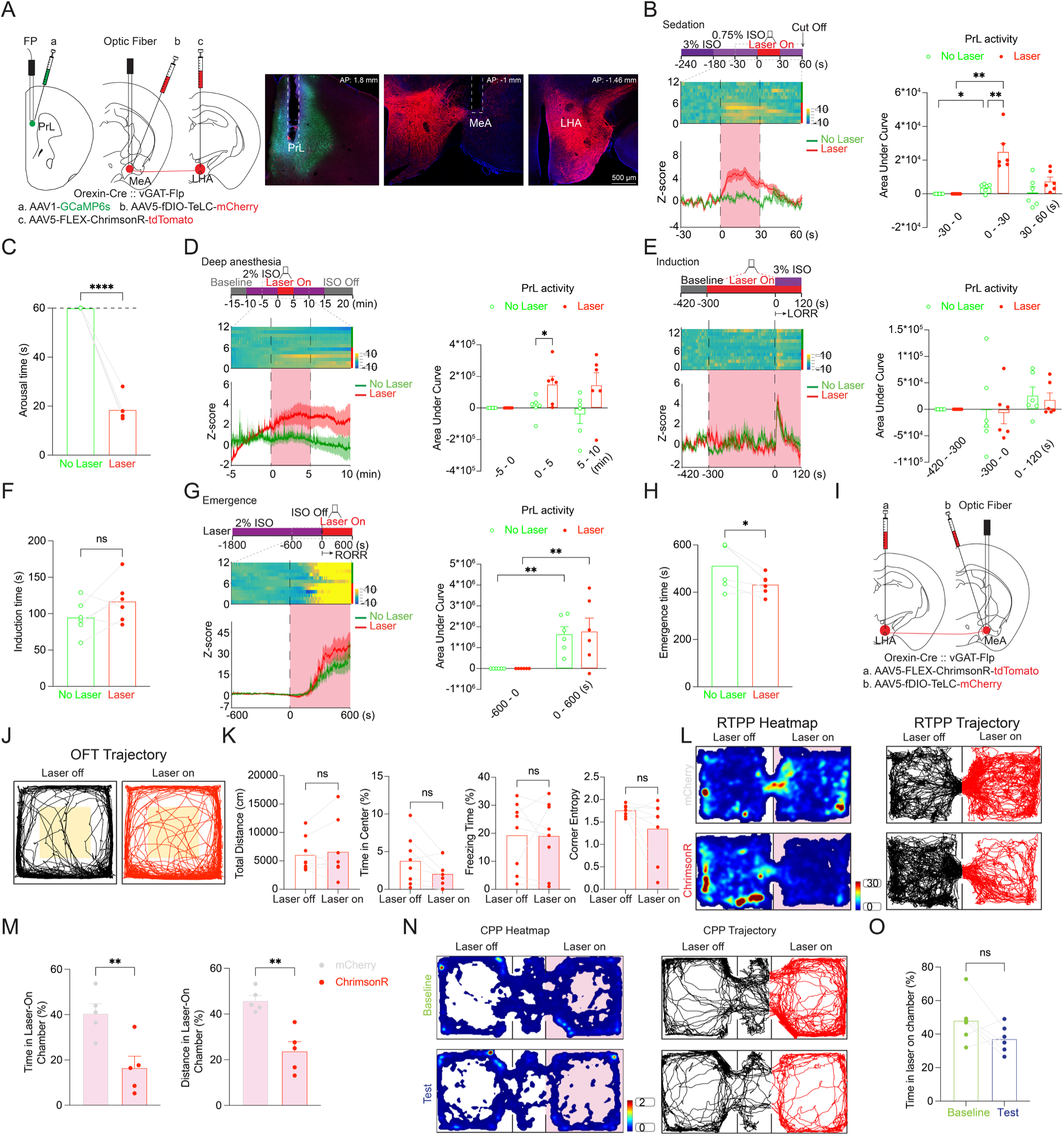
MeA^vGAT^ output gates LHA^Ox^→MeA-driven cortical activation, arousal, and valence-related behavior. (**A**) Intersectional strategy in Orexin-Cre::vGAT-Flp mice to stimulate LHA^Ox^ terminals (Cre-dependent ChrimsonR in LHA^Ox^) while blocking MeA^vGAT^ synaptic output (Flp-dependent TeLC in MeA^vGAT^) and monitoring prelimbic cortex (PrL) activity (GCaMP6s in PrL); optic fibers were implanted above the MeA and PrL. Representative images confirm the expression of ChrimsonR, TeLC, and GCaMP6s. (**B**, **D**) PrL activity during LHA^Ox^→MeA stimulation with MeA^vGAT^ output silenced under isoflurane sedation (**B**) and 2% isoflurane anesthesia (**D**) with quantification. (**C**) Despite MeA^vGAT^ output silencing, LHA^Ox^→MeA stimulation induces arousal from light anesthesia. (**E** to **F**) During induction, pre-activation of LHA^Ox^→MeA terminals with MeA^vGAT^ output silenced does not modulate PrL activity (**E**) or delay loss of the righting reflex (LoRR) (**F**). (**G** to **H**) During emergence, stimulation does not increase PrL activity (**G**) but accelerates recovery of the righting reflex (RoRR) (**H**). (**I**) Strategy for behavioral assays: viral expression and optic fiber implantation. (**J** to **K**) Representative open-field trajectories (**J**) and quantification of exploration and defensive metrics (**K**) during LHA^Ox^→MeA stimulation with MeA^vGAT^ output silenced. (**L** to **M**) Representative RTPP heatmaps/trajectories (**L**) and quantification showing reduced time spent and distance traveled in the laser-paired chamber in ChrimsonR mice relative to controls under MeA^vGAT^ output silencing (**M**). (**N** to **O**) Representative CPP heatmaps/trajectories during baseline and test sessions (**N**) and quantification showing no conditioned change in time spent in the stimulation-paired chamber (**O**). Data are mean ± S.E.M.; statistical details and exact n values are provided in Supplementary Table 1. *p ≤ 0.05, **p ≤ 0.01, ***p < 0.001, ****p < 0.0001.

Under isoflurane sedation, LHA^Ox^→MeA stimulation still increased PrL activity and induced behavioral arousal (Fig. 6B–C), and under deep anesthesia (2% isoflurane), it elevated PrL activity during the early post-stimulation window (Fig. 6D). In contrast, stimulation no longer altered PrL activity or LoRR during induction (Fig. 6E–F), and no longer increased PrL activity during emergence (Fig. 6G). While still significantly shortened RoRR latency (Fig. 6H). These results indicate that MeA^vGAT^ synaptic output is required for orexin-dependent modulation of induction and for state-dependent cortical engagement, but is dispensable for orexin-driven arousal from light anesthesia or accelerated emergence.

Finally, we asked whether MeA^vGAT^ output is required for LHA^Ox^→MeA-driven vigilance-like behavioral bias by activating the LHA^Ox^→MeA projection while silencing MeA^vGAT^ output (Fig. 6I–O). Under these conditions, LHA^Ox^→MeA stimulation no longer suppressed exploration-related measures in the open field (Fig. 6K). In RTPP, following blockade of the MeA^vGAT^ synaptic output, LHA^Ox^→MeA stimulation decreased both time spent and distance traveled in the laser-paired compartment, indicating a shift from preference to avoidance (Fig. 6M). CPP likewise showed no conditioned preference shift (Fig. 6O). Together, these findings indicate that MeA^vGAT^ synaptic output is required for LHA^Ox^→MeA-driven vigilance-like behavioral bias and real-time valence effects.

## Discussion

General anesthesia, like natural sleep, produces a reversible loss of consciousness and engages sleep–wake circuits (5, 6, 51, 52). Despite extensive work on arousal circuits, the limbic pathways that couple anesthetic arousal to vigilance-like behavioral states remain poorly defined. The present study addresses this gap by identifying a lateral hypothalamic orexin to medial amygdala (LHA^Ox^→MeA) circuit that links anesthetic state transitions to vigilance-like behavioral bias.

### Medial amygdala as a projection-specific node for orexin function

We identify the MeA as a projection-defined orexin target that links arousal control to behavioral bias: the LHA^Ox^→MeA projection forms functional synapses and promotes arousal from NREM sleep and anesthesia (Fig. 1; Supplementary Figs. 1 to 2). Unlike global LHA^Ox^ activation–associated analgesia (22, 37–39), selective LHA^Ox^→MeA activation did not increase thermal pain tolerance, supporting projection-specific dissociation of orexin functions (Supplementary Fig. 2D).

Orexin–amygdala interactions have been studied primarily in the context of fear, stress, and emotional learning, with orexin acting both indirectly via ascending arousal systems and directly within amygdala nuclei (53–55). Early anatomical mapping established dense hypocretin/orexin innervation across amygdala subregions (56), and receptor mapping confirmed amygdala expression of Ox1R and Ox2R (57). Functionally, much of the mechanistic literature has emphasized central and basolateral amygdala circuits in shaping anxiety- and fear-related behaviors, including orexin-dependent modulation of amygdala excitability and defensive responding (58). In contrast, comparatively little is known about how orexin engages the MeA to couple arousal state control to vigilance-like behavioral strategy selection. Our findings on the LHA^Ox^→MeA coupling orexin signaling to anesthetic state transitions and acute vigilance bias extend orexin–amygdala biology beyond canonical fear circuitry.

### State-dependent orexin control of induction, arousability, and emergence

Our results also refine the view that induction, steady-state sedation, and emergence represent distinct control regimes rather than equivalent manifestations of a single arousal process (42). We therefore examined all three phases to distinguish effects on transition thresholds (entry into and exit from unconsciousness) from within-state arousability under a fixed anesthetic drive.

Orexin has been implicated in facilitating emergence with limited effects on induction (41, 42). One possibility is that peptide-dependent contributions during induction are constrained by the rapid timescale of the transition unless orexin tone is elevated in advance. Consistent with this idea, pre-activation of the LHA^Ox^→MeA projection increased MeA orexin release and delayed loss of righting reflex during induction (Fig. 2N–O). In contrast, despite robust Jaws-mediated suppression of orexin excitability ex vivo, pre-inhibition did not measurably reduce MeA orexin signal in the awake baseline and did not shorten induction time (Supplementary Fig. 3), potentially reflecting low baseline orexin tone during the light phase (59, 60). Once induction began, inhibition accelerated the decline in orexin signal (Supplementary Fig. 3D), consistent with an interaction between anesthetic drive and reduced orexin tone during the transition into unconsciousness. Given isoflurane’s broad molecular targets (61–63), this pattern likely reflects convergent suppression of arousal-promoting processes rather than a one-to-one mapping between orexin release and induction timing.

Emergence has immediate clinical relevance: delayed awakening can increase the risk of airway obstruction and hypoxemia, consume substantial clinical resources, and prompt costly neuroradiologic evaluations (64). In our study, activation of the LHA^Ox^→MeA projection augmented MeA orexin release and accelerated emergence from isoflurane anesthesia (Fig. 2P–Q). Inhibiting the LHA^Ox^→MeA projection did not slow orexin signal recovery or prolong emergence (Supplementary Fig. 3F–G), suggesting redundancy across orexin projections and/or compensatory arousal circuits that preserve behavioral recovery when a single projection is suppressed.

### MeA^vGAT^ output gates orexin-driven cortical engagement

At the circuit level, RNAscope mapping and cell-type-specific photometry identify MeA^vGAT^ neurons as a principal downstream substrate engaged by LHA^Ox^ input (Fig. 4; Supplementary Figs. 4–5). Prior work has shown that orexin neurons can stabilize arousal by recruiting subcortical circuitry that suppresses sleep-promoting nodes and facilitates rapid state switching (65, 66). Consistent with this framework, LHA^Ox^→MeA activation increased MeA^vGAT^ Ca²⁺ signals across sedation, deep anesthesia, and emergence (Fig. 4), positioning this inhibitory population as a state-spanning node linking orexin input to anesthetic transitions. During induction, pre-activation elevated MeA^vGAT^ activity relative to time-matched controls before isoflurane onset, but this difference diminished after anesthetic exposure (Fig. 4G). Importantly, LoRR was still delayed (Fig. 4H), suggesting that induction latency can be influenced by transient pre-onset dynamics, recruitment of a sparse vGAT subpopulation, synaptic output changes not captured by population-averaged photometry, or compression of measurable dynamics under deepening anesthesia.

Although MeA^vGAT^ neurons are inhibitory, activation of the inhibitory population can promote wakefulness by suppressing sleep-promoting nodes and facilitating state transitions (67–69). Consistently, MeA^vGAT^ activation is sufficient to drive PrL cortical engagement and promote arousal and emergence (Fig.5). Moreover, across manipulations, orexin terminal activation (Fig.4G–H) and direct activation of MeA^vGAT^ neurons (Fig.5E–F) prolonged LoRR, whereas silencing MeA^vGAT^ synaptic output abolished this induction prolongation (Fig.6E–F). Together, these data support a state-dependent gating model in which MeA^vGAT^ synaptic output is required for orexin-dependent modulation of the induction and for state-dependent cortical engagement, while orexin-driven arousal from light anesthesia and accelerated emergence can be preserved when MeA^vGAT^ output is blocked. In addition, under deep anesthesia, direct activation of MeA^vGAT^ neurons was insufficient to increase PrL activity (Fig.5D), whereas LHA^Ox^→MeA activation still produced a detectable PrL increase when MeA^vGAT^ output was silenced (Fig.6D). Together, these findings indicate that orexin terminal activation can influence cortical activity through additional pathways beyond MeA^vGAT^ synaptic output. Defining these parallel routes and their state dependence will be important for understanding how orexin inputs are distributed across distinct arousal-control channels.

### Coupling arousal to vigilance-like behavioral bias

Beyond anesthesia state control, our data link LHA^Ox^→MeA signaling to vigilance-like behavioral bias. The amygdala coordinates defensive vigilance and strategy selection during heightened arousal, and the posteroventral medial amygdala (MePV) glutamatergic neurons have been implicated in coupling wakefulness to anxiety-like phenotypes (29, 70–72). In our assays, LHA^Ox^→MeA activation produced an acute defensive vigilance-like state rather than freezing. It reduced open field exploration and center engagement without increased freezing, and a shift to stereotyped, corner-focused locomotion (Fig.3). In RTPP, stimulation increased time and distance in the laser-paired chamber, but movement was corner-confined and repetitive, consistent with localized “scanning” rather than exploration (Fig.3). Together, these findings support a model in which orexin engagement of MeA circuitry shapes both arousal state and the defensive behavioral mode expressed upon arousal.

Direct activation of MeA^vGAT^ neurons induced real-time place preference and drove perimeter-biased, distributed locomotion consistent with thigmotactic “patrolling”, a distinct vigilance-like pattern from the corner-confined pattern observed during LHA^Ox^→MeA activation (Fig.5). Prior work indicated that MeA^vGAT^ output can alleviate post-shock freezing and anxiety-like behaviors in specific contexts (73), consistent with a role in selecting defensive strategies rather than uniformly promoting immobility. Notably, silencing MeA^vGAT^ output altered the behavioral impact of LHA^Ox^→MeA activation in a task-dependent manner: in the OFT, stimulation no longer reduced exploration-related measures (total distance, center time, freezing time, corner entropy), whereas in the RTPP, silencing converted orexin-driven real-time preference into avoidance, indicating MeA^vGAT^ synaptic output is required for orexin-driven real-time preference (Fig.6). CPP remained negative across conditions, consistent with acute and state-dependent effects rather than durable conditioned reinforcement.

This study opens several avenues for deeper mechanistic resolution. First, establishing receptor dependence will require MeA-restricted loss-of-function of Ox1R and Ox2R during anesthetic transitions and vigilance-like assays. Second, because orexin neurons co-release additional transmitters, including glutamate and dynorphin, co-transmission may contribute to the circuit effects of terminal stimulation (74–76). Third, because fiber photometry reports population-averaged signals, sparse subpopulations, output-defined ensembles, or synaptic dynamics not captured by bulk Ca²⁺ signals may be especially important for gating anesthesia state-transitions. Finally, defining the downstream targets of MeA^vGAT^ output will be essential for mechanistic closure and will test whether this limbic node acts through intermediate subcortical relays and/or cortical inhibitory networks to translate orexin drive into cortical engagement and state-dependent behavioral bias.

Overall, the present study identified an LHA^Ox^→MeA projection that links orexin-dependent control of anesthetic state transitions with vigilance-like behavioral bias (Fig. 7). It provides a circuit-level entry point for understanding how perioperative arousal can be coupled to threat-sensitive behavioral states and implicates MeA^vGAT^ as a key gate shaping both anesthetic transitions and vigilance-related phenotypes.

**Fig. 7.**
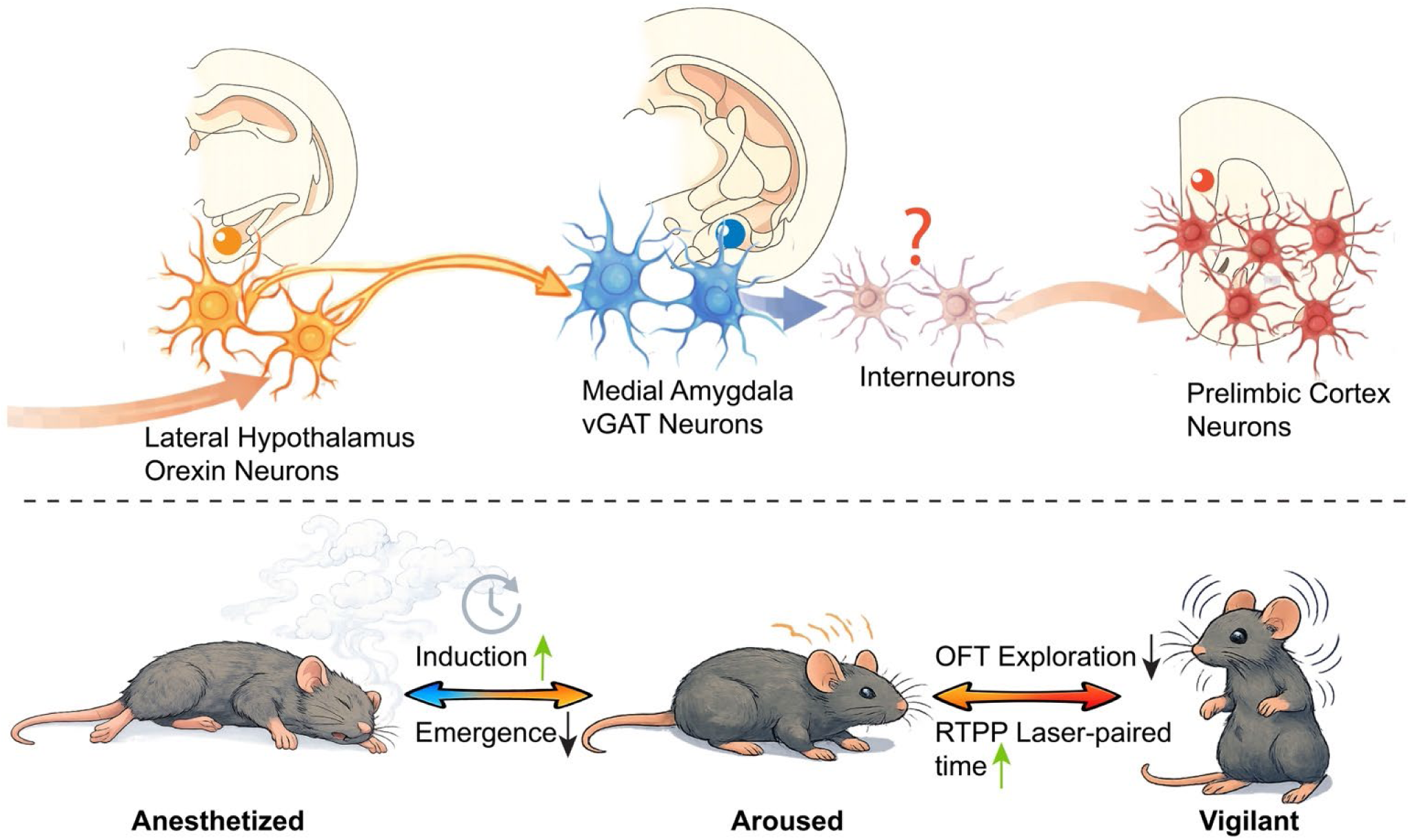
An LHA^Ox^→MeA^vGAT^ pathway links orexin-dependent state transitions to vigilant behavioral bias. Lateral hypothalamic orexin (LHA^Ox^) neurons project to the medial amygdala (MeA) and engage MeA^vGAT^ neurons through predominantly Ox2R-mediated signaling. Activation of the LHA^Ox^→MeA pathway increases local orexin release, promotes arousal from isoflurane sedation, delays induction (LoRR), and accelerates emergence (RoRR). MeA^vGAT^ neurons represent a key downstream substrate: their activation is sufficient to drive prefrontal cortical engagement and bias vigilance-like behavior, whereas blocking MeA^vGAT^ synaptic output disrupts orexin-dependent induction delay and flips real-time place preference effects. The downstream targets linking MeA^vGAT^ output to the cortex may involve interneuron populations.

## Materials and Methods

A total of 144 male mice (6–20 weeks old; 18–36 g) were used. Mouse lines included Orexin-Cre-2A-EGFP (Orexin-Cre; n = 82), vGAT-2A-FlpO-Δ (vGAT-Flp; n = 18), Orexin-Cre::vGAT-Flp (n = 31), and Orexin-Cre::vGlut2-Flp (n = 13). The Orexin-Cre line was a gift from Dr. Akihiro Yamanaka (Nagoya University, Japan) and was maintained on a C57BL/6 background. C57BL/6J (stock no. 000664), Vglut2-IRES2-FLPo-D (Vglut2-Flp; stock no. 030212), and vGAT-2A-FlpO-Δ (vGAT-Flp; stock no. 029591) mice were obtained from The Jackson Laboratory. Orexin-Cre::vGlut2-Flp and Orexin-Cre::vGAT-Flp mice were generated by crossing male Vglut2-Flp or vGAT-Flp mice with female Orexin-Cre mice. Only male offspring positive for both Cre and Flp, confirmed by genotyping, were used in experiments.

Mice were housed under controlled conditions with ad libitum access to food and water on a 12-hour light/dark cycle (lights on 07:00, lights off 19:00) at 20–22°C. Only male mice were used to avoid estrous-cycle–associated hormonal variation; additionally, because the Orexin-Cre driver is X-linked in this line, male mice provide consistent Cre dosage. All procedures were approved by the Institutional Animal Care and Use Committee at the University of California, San Francisco.

### Virus Injection

#### Viral vectors

AAV vectors were obtained from Addgene unless otherwise noted. Viruses used included Cre-dependent synaptic labeling (AAV1-FLEX-tdTomato-T2A-SypEGFP), Cre-dependent optogenetic actuators (AAV5-FLEX-ChrimsonR-tdTomato; AAVrg-DIO-ChR2-mCherry; AAV5-FLEX-Jaws-GFP; AAV5-DIO-mCherry; AAVrg-DIO-mCherry), Flp-dependent reporters/actuators (AAV1-fDIO-GCaMP6f; AAV8-FLPX-ChrimsonR-tdTomato), and AAV1-GCaMP6s for cortical photometry.

For orexin release measurements, the OxLight sensor plasmid (AAV2-OxLight; Addgene 169792) was obtained from Addgene, and virus was produced in-house following the Salk Institute Viral Vector Core protocol. A Flp-dependent tetanus toxin light chain construct (pAAV-hSyn-fDIO-TeLC-mCherry) was generated using one-step cloning (ClonExpress Ultra; Vazyme). The TeLC fragment was PCR-amplified from pAAV-hSyn-FLEX-TeLC-P2A-dTomato (Addgene 159102) and substituted for hM4D(Gi) in pAAV-hSyn-fDIO-hM4D(Gi)-mCherry (Addgene 154867), yielding an hSyn-driven, Flp-dependent TeLC-mCherry cassette. Recombinant AAV5-hSyn-fDIO-TeLC-mCherry was packaged by the HHMI Janelia Viral Core. All viruses were aliquoted and stored at −80°C; aliquots were single-use (no refreezing after thaw).

### Stereotaxic surgery and injection parameters

Intracranial injections were performed using a robotic stereotaxic system (Neurostar, Germany) under balanced anesthesia. Mice received isoflurane (Henry Schein Animal Health) as needed during surgery, and perioperative analgesia/antibiotic coverage (meloxicam 5 mg/kg s.c.; buprenorphine 0.1 mg/kg s.c.; ampicillin 10–20 mg/kg s.c.; local bupivacaine 0.25%, 25 µl s.c.). Viruses were delivered using a 2 µl Hamilton Neuros syringe (7000 series) with a 0.5 µl injection volume per site infused over 10 min. The needle was left in place for 5 min, then slowly withdrawn. The incision was closed using wound clips (BD Autoclip).

### Injection coordinates

Coordinates (mm) are relative to bregma:

Hypothalamus (HY/LHA): AP −1.0; ML ±0.95; DV 5.0–5.1

Medial amygdala (MeA): AP −1.0; ML ±2.0; DV 4.9–5.0

Diagonal band nucleus (NDB): AP +1.0; ML 0.0; DV 4.7–4.8

Prelimbic cortex (PrL): AP +1.8; ML ±0.3; DV 2.3–2.4

### Fiber Implantation

For optogenetic stimulation and EEG/EMG recordings, mice were implanted with bilateral optic fibers targeting the MeA (400-µm core, 0.39 NA; RWD Life Science; R-FOC-BL400C-39NA). Fiber tips were placed at AP −1.0 mm, ML ±2.0 mm, DV −5.0 mm (relative to bregma).

For fiber photometry, mice were implanted with unilateral optic fibers (200-µm core, 0.39 NA; RWD Life Science; R-FOC-BL200C-39NA) targeting either the left or right MeA (AP −1.0 mm, ML ±2.0 mm, DV −5.0 mm). For experiments monitoring prefrontal activity, an additional unilateral fiber was implanted above the prelimbic cortex (PrL) (AP +1.8 mm, ML ±0.3 mm, DV −2.4 mm).

### EEG/EMG Implantation

During the same surgical session as viral injection and optic fiber implantation, mice were implanted with an EEG/EMG headmount (8201, Pinnacle Technologies). Four skull holes were drilled using a 23-gauge needle at two frontal sites (AP +1.0 mm, ML ±1.25 mm) and two parietal sites (AP −3.0 mm, ML ±2.5 mm). Four stainless-steel screws with attached leads (8493, Pinnacle Technologies) were inserted into the skull and connected to a six-pin EEG/EMG headmount via soldered leads. EMG leads were inserted into the nuchal (neck) musculature. The assembly was secured with dental cement (C&B Metabond Quick Adhesive Cement System, Parkell). Animals were allowed to recover for 4 weeks before behavioral and physiological experiments.

### EEG recording with optogenetic manipulation

Mice were briefly anesthetized in an induction chamber with 3% isoflurane for 1 min and then connected to the EEG/EMG acquisition system (8200-K1-SL2, Pinnacle Technologies). Bilateral implanted optic fibers were coupled via ceramic mating sleeves (R-MS-1.25, RWD Life Science) to a bifurcated patch cable (400-µm core, 0.39 NA; FC/PC to 1.25-mm ferrules; 1 m; Thorlabs BFYL4LF01), which was connected to a 473-nm laser (Aurora-200, Newdoon). Mice were acclimated to the recording setup for 24 h before EEG experiments; on each recording day, mice were additionally acclimated for 30 min before data collection.

### Fiber photometry recording with optogenetic manipulation

Mice were briefly anesthetized in an induction chamber with 3% isoflurane for 1 min and connected to a multichannel fiber photometry system (R821, RWD Life Science). A low-autofluorescence patch cord (Doric Lenses; MFP_400/440/3000-0.37_1m_FCM-MF1.25(F)_LAF) was coupled to the implanted fiber(s). GCaMP signals were excited at 470 nm, and an isosbestic 410-nm channel was collected as a reference for motion/photobleaching correction. For simultaneous optogenetic manipulation, a 635-nm laser (IOS-635, RWD Life Science) was integrated with the R821 system to stimulate ChrimsonR-expressing terminals while recording fluorescence. Mice were acclimated to the recording setup for ≥30 min before data collection. Optogenetic stimulation parameters are described in the corresponding behavioral and anesthesia paradigms.

### Immunohistochemistry

Mice were transcardially perfused 4–8 weeks after viral injection with ice-cold PBS followed by 4% paraformaldehyde (PFA) in PBS (Electron Microscopy Sciences). Brains were post-fixed in 4% PFA for 6 h at 4°C and cryoprotected in 30% sucrose for ≥2 days until saturated. Coronal or sagittal sections (35 µm) were cut on a cryostat (Leica CM3050S).

Free-floating sections were blocked for 1 h at room temperature in PBS containing 5% donkey serum, 3% bovine serum albumin (BSA), and 0.3% Triton X-100, then incubated with primary antibodies at 4°C for 16–24 h. Sections were washed 3× in PBS (10 min each) and incubated with secondary antibodies for 2 h at room temperature, followed by 3× PBS washes. Sections were mounted with DAPI Fluoromount-G (SouthernBiotech, 0100-20).

Primary and secondary antibodies (diluted in blocking solution) were as follows: mCherry: chicken anti-mCherry (1:400; Novus Biologicals, NBP2-25158); Cy3-conjugated goat anti-chicken IgY (1:400; Jackson ImmunoResearch, 103-165-155). cFos: rabbit anti-cFos (1:100; Cell Signaling Technology, 2250S); Alexa Fluor 488–conjugated donkey anti-rabbit IgG (1:400; Jackson ImmunoResearch, 711-546-152).

### Imaging

Confocal images were acquired on a Leica TCS SP8 confocal microscope. Tile-scan imaging was performed using Leica Application Suite software with automated tiling settings.

### Behavioral Tests

All behavioral and anesthesia-transition experiments were conducted between 09:00 and 18:00 with age-matched cohorts. For anesthesia procedures, chambers were placed on a temperature-controlled heating pad to maintain body temperature at 35–37°C. Throughout the manuscript, the term “laser-paired chamber/compartment” refers to the side in which optical stimulation was delivered (closed-loop for RTPP; designated stimulation side for CPP). Behavioral videos and MATLAB-based analyses were performed using animal IDs with the experimenter blinded to group and stimulation condition until completion of quantification.

### Isoflurane arousal from sedation

Mice undergoing optogenetic manipulation were induced with 3% isoflurane for 1 min and then connected to the recording/stimulation apparatus. Animals were transferred to a sealed clear chamber with optic fibers routed through a port and acclimated for 30 min. The chamber was equilibrated with 3% isoflurane for 1 min, switched to 0.75% isoflurane for 3 min, and then optical stimulation (473 or 635 nm) was delivered for 30 s. Isoflurane was maintained for an additional 1 min after stimulation and then replaced with 100% O₂ for recovery. Arousal latency was defined as the time from laser onset to the first observable signs of arousal (body movement, kicking, or tail elevation). Each mouse completed three trials in a single day; control mice received identical procedures.

### Isoflurane induction

Mice were connected to the fiber photometry system and placed in a sealed clear chamber. Baseline photometry was recorded for 10 min. Optical stimulation (635 nm; 20 Hz; 20-ms pulse width) began 5 min before induction and continued through the end of the trial. Isoflurane (3%) was then delivered for 3 min. Loss of righting reflex (LoRR) latency was defined as the time from isoflurane onset to loss of the righting reflex. Each mouse underwent two sessions separated by 1 day.

### Isoflurane emergence

Mice were anesthetized with 2% isoflurane for 30 min and then maintained in the same chamber under 100% O₂ with either optical stimulation (20 Hz; 20-ms pulse width) or no stimulation. Recovery of righting reflex (RoRR) was defined as the mouse turning over and standing on all four paws; emergence latency was measured from isoflurane discontinuation to RoRR. Sessions were repeated three times at 3-day intervals; controls received identical procedures.

### Hot plate test

Mice were connected to optic fibers and received optical stimulation or no stimulation for 5 min (20 Hz; 20-ms pulse width; 1 s on/1 s off), followed immediately by the hot plate assay. Mice were placed in an acrylic enclosure on a 55°C hot plate and stimulation continued throughout the test. Paw withdrawal latency was defined as the first nocifensive response (paw licking, fanning, or jumping), at which point mice were removed. A cutoff of 45 s was applied. Trials were performed three times in one day (09:00–15:00) with 60-min intertrial intervals and repeated twice more at 3-day intervals; controls received identical procedures.

### Open field test (OFT)

Mice were placed in a corner of a white open-field arena (60 × 60 × 40 cm) and allowed to explore freely. Behavior was video recorded for 20 min. The center zone was defined as a 30 × 30 cm square. Optical stimulation (635 nm; 20 Hz; 20-ms pulse width) was delivered continuously throughout the 20-minute recording period. The arena was cleaned with 75% ethanol between trials. Trajectories, total distance traveled, center time, freezing time, and corner entropy were quantified offline in MATLAB.

### Real-time place preference (RTPP)

A transparent chamber (40 × 20 × 20 cm) was divided into two equal compartments. Mice were placed initially in the non-stimulation compartment and allowed to explore for 20 min. Optical stimulation (635 nm; 20 Hz; 20-ms pulse width) was delivered in a closed-loop manner when the animal entered the stimulation compartment and terminated upon exit. Compartment assignment was randomized and counterbalanced across mice. Behavior was video recorded and position tracking/closed-loop laser control was implemented via an RWD photometry system (R821). Heatmaps, trajectories, time spent, and distance traveled in the stimulation-paired compartment were analyzed offline in MATLAB.

### Conditioned place preference (CPP)

A transparent chamber (60 × 30 × 40 cm) comprised a middle compartment (10 × 30 × 40 cm) and two side compartments (25 × 30 × 40 cm) with distinct visual and tactile cues. The stimulation-paired side was randomized and counterbalanced. Mice underwent 3 days of habituation (30 min/day; free exploration; no stimulation) followed by a 20-min baseline test. Conditioning consisted of 3 days, with two 20-minute sessions/day (morning and afternoon; ≥6 h apart) with doors closed. During the stimulation session, mice were placed in the assigned compartment and received optical stimulation (635 nm; 20 Hz; 20-ms pulse width); no stimulation was delivered during the alternate session. On the test day, doors were removed, and mice freely explored for 20 min, as in baseline. Behavior was video-recorded and position tracking/laser control was implemented via RWD R821. Heatmaps, trajectories, and time spent in the stimulation-paired compartment were analyzed in MATLAB.

### Electrophysiology

Acute brain slices were prepared from 8–12-week-old Orexin-Cre mice. Mice were anesthetized with isoflurane and transcardially perfused with ice-cold, carbogenated (95% O₂/5% CO₂) choline-based cutting solution containing (in mM): 89.5 choline chloride, 20.6 Tris, 2.5 KCl, 1.2 NaH₂PO₄·2H₂O, 20 HEPES, 5 sodium ascorbate, 2 thiourea, 3 sodium pyruvate, 25 glucose, 30 NaHCO₃, 10 MgSO₄·7H₂O, 0.5 CaCl₂·2H₂O (300–310 mOsm; pH 7.4). Brains were removed and sectioned at 250 µm using a vibratome (Leica VT1000S). Slices recovered for 10–15 min at 34°C in cutting solution, then were transferred to a HEPES-based holding aCSF containing (in mM): 92 NaCl, 2.5 KCl, 1.25 NaH₂PO₄, 30 NaHCO₃, 20 HEPES, 25 glucose, 2 thiourea, 5 Na-ascorbate, 3 Na-pyruvate, 2 CaCl₂·4H₂O, 2 MgSO₄·7H₂O (300–310 mOsm; pH 7.3–7.4) for 45–60 min at room temperature. For recordings, slices were superfused (3 ml/min) with carbogenated recording aCSF containing (in mM): 124 NaCl, 2.5 KCl, 1.2 NaH₂PO₄·2H₂O, 24 NaHCO₃, 5 HEPES, 12.5 glucose, 2 CaCl₂·2H₂O, 2 MgSO₄·7H₂O (300–310 mOsm; pH 7.3–7.4 adjusted with NaOH).

Patch electrodes (3–5 MΩ) were pulled from borosilicate glass (BF150-86-10, Sutter) and filled with a potassium-based internal solution containing (in mM): 145 K-gluconate, 10 HEPES, 1 EGTA, 2 Mg-ATP, 0.3 Na₂-GTP, 2 MgCl₂ (pH 7.2–7.3 with KOH; 290–300 mOsm). Neurons were visualized with a 60× water-immersion objective (Zeiss). Whole-cell recordings were acquired using a MultiClamp 700B amplifier with Digidata 1322A digitizer and pClamp (Molecular Devices), sampled at 10 kHz, and analyzed in Clampfit.

Optogenetic stimulation was delivered with a 635-nm laser (IOS-635, RWD Life Science) using 20-Hz, 20-ms pulses (five trials per neuron). Cells were voltage-clamped at −65 mV to record EPSCs and at 0 mV to record IPSCs; current-clamp recordings were used to assess action potential firing. Orexin receptor antagonists were bath-applied as indicated: TCS OX2 29 (30 µM; MilliporeSigma SML2879) and SB-334867 (10 µM; MilliporeSigma SML1530).

### RNA-scope in situ hybridization

Adult C57BL/6J mice (14–16 weeks) were transcardially perfused as described in the Immunohistochemistry section. Brains were post-fixed in 4% PFA at 4°C for 24 h, cryoprotected, and sectioned at 15 µm. Sections were mounted on slides and stored at −80°C until processing. Quantification was performed within the medial amygdala using anatomically defined boundaries based on a standard mouse brain atlas (to be specified), with the analysis conducted blinded to experimental condition.

RNAscope multiplex fluorescent in situ hybridization was performed using the RNAscope Multiplex Fluorescent Detection Reagents v2 kit (Advanced Cell Diagnostics, 323110) following the manufacturer’s protocol. Probes were obtained from Advanced Cell Diagnostics: Ox1R (466631-C1), Ox2R (581631-C3), vGAT (319191-C2), vGlut2 (428871-C2), and TH (317621-C2).

Briefly, slides were post-fixed in 4% PFA for 2 h, washed in PBS, treated with H₂O₂ for 10 min at room temperature, rinsed in DEPC-treated water, and subjected to target retrieval (98–102°C for 5 min). After rinsing and dehydration in 100% ethanol, slides were dried at 60°C for 5 min, rehydrated, and incubated with Protease III for 15 min at 40°C in a HybEZ oven (ACD). Probe mixtures were hybridized for 2 h at 40°C, followed by signal amplification (AMP1, 30 min; AMP2, 30 min; AMP3, 15 min). Channel development was performed sequentially using HRP-based amplification with Opal fluorophores: Opal 520 (Akoya, FP1487001KT), Opal 620 (Akoya, FP1495001KT), and Opal 690 (Akoya, FP1497001KT), with HRP blocking steps between channels. Slides were washed with ACD wash buffer between each step. Sections were mounted with DAPI Fluoromount-G (SouthernBiotech, 0100-20) and coverslipped.

Images were acquired on a Leica TCS SP8 confocal microscope using a 40× oil-immersion objective. Image contrast/levels were adjusted uniformly in FIJI/ImageJ. Quantification was performed in QuPath v0.5.1 using automated detection pipelines.

### Statistics

Data were analyzed by experimenters blinded to the experimental condition. Statistical tests were selected based on experimental design: paired two-tailed t tests were used for within-subject comparisons, unpaired two-tailed t tests (with Welch’s correction when variances were unequal) for between-group comparisons, and one-way or two-way ANOVA for experiments with one or two factors, respectively, followed by multiple-comparisons procedures as specified in the figure legends. Data are presented as mean ± S.E.M. Statistical significance was defined as P < 0.05. Analyses were performed in GraphPad Prism 10.6.1 (GraphPad Software, San Diego, CA).

## Acknowledgments

We thank Drs. Lily Jan and Yuh-Nung Jan for sharing lab resources; Dr. Akihiro Yamanaka for sharing the frozen embryo of the orexin-Cre mouse line; Dr. Wendy W.S. Yue for insightful discussions and proofreading; Tong Cheng and Marena Tynan-La Fontaine for their technical support. All experimental data were generated at the University of California, San Francisco.

## Author contributions

W.Z. and X.X. designed and performed research; X.X. drafted the manuscript; C.C. contributed to AAV5-hSyn-fDIO-TeLC-mCherry plasmid reconstruction, AAV2-Oxlight virus production, and RNAscope experimental design; W.Z. contributed to the project administration, manuscript revision, and funding acquisition.

## Data and materials availability

All data generated in this study are included within the main text or supplementary materials, with source data provided. This includes individual data points and averaged values presented in both the figures and supplementary information. The corresponding authors can provide raw data upon request.

