## Supplementary Data for "Hypothalamic Orexin Input to the Medial Amygdala Links Vigilance to Arousal"

### Supplementary table.1

| Figure | Data | Sample size | Statistical test | Exact p value |
|---|---|---|---|---|
| Fig. 1. F<br>Amplitudes of EPSCs | 35.16 ± 3.28 pA | n=35 | Two-tailed paired t-test, t = 10.71, df = 34. | p < 0.0001 |
| Fig. 1. F<br>Amplitudes of IPSCs | 67.81 ± 9.85 pA, | n=17 | Two-tailed paired t-test, t = 6.886, df = 16. | p < 0.0001 |
| Fig. 1. F<br>Latency | EPSCs: 10.06 ± 0.89 ms,<br>IPSCs: 10.48 ± 0.81 ms; | EPSCs: n=35,<br>IPSCs: n=17 | Welch's t test, t = 0.3459, df = 46.47. | p = 0.731 |
| Fig. 1. I | $Ox1R^+$ (3.05 ± 0.51), $Ox2R^+$ (38.13 ± 2.39), and $Ox1R^+$ + $Ox2R^+$ (14.88 ± 4.31); | n = 8 slices from 4 brains | Ordinary one-way ANOVA, F(2, 21) = 38.94, p < 0.0001; Šídák's multiple-comparisons test. | Ox1R only vs Ox2R only: p < 0.0001, Ox1R only vs Ox1R + Ox2R: P = 0.0241, Ox2R only vs Ox1R + Ox2R: p < 0.0001. |
| Fig. 1. K<br>EPSCs amplitude with Ox1R antagonist SB-334867 | Before, 30.27 ± 3.1; SB-334867, 9 ± 3.35; Washout, 29.73 ± 2.03; | n = 15; | Repeated measures one-way ANOVA, F(1.274, 17.84) = 18.37, p = 0.0002, Šídák's multiple comparisons. | baseline vs SB-334867: P = 0.0021, SB-334867 vs washout: p = 0.0011. |
| Fig. 1. K<br>EPSCs amplitude with Ox2R antagonist TCS Ox2 29 | Before, 31.98 ± 3.09; TCS Ox2 29, 3.82 ± 1.84; Washout, 28.61 ± 2.84; | n=19. | Repeated measures one-way ANOVA, F(1.891, 34.04) = 48.74, p < 0.0001, Šídák's multiple comparisons. | baseline vs TCS OX2 29: p < 0.0001, TCS OX2 vs washout: p < 0.0001. |

| | | | | |
|---|---|---|---|---|
| Fig. 2. D | mCherry control group： 1.95 ± 0.36;<br>ChR2 group: 7.58 ± 0.72 | mCherry, n = 6; ChR2, n = 7 | Welch's t test; t = 7.005, df = 8.709. | p < 0.0001 |
| Fig.2G<br>EEG power spectral density in the mCherry group | Delta: pre, 45.65 ± 2.97; stim, 43.17 ± 3.33;<br>Theta: pre, 24.71 ± 1.72; stim, 24 ± 1.48;<br>Alpha: pre, 12.1 ± 1.35; stim, 12.82 ± 1.98 ;<br>Beta: pre, 17.33 ± 1.03; stim, 19.78 ± 1.26; | n=10 | Two-way ANOVA, F(1, 9) = 0.0239, p = 0.8805; Šidák's multiple comparison test. | Pre vs Stim: Delta, p = 0.0896;<br>Theta, p = 0.9721;<br>Alpha, p = 0.9107;<br>Beta: p = 0.077. |
| Fig.2G<br>EEG power spectral density in ChR2 group | Delta: pre, 45.3 ± 2.87; stim: 37.73 ± 2.24;<br>Theta: pre, 27.13 ± 2.01; stim, 25.03 ± 2.4;<br>Alpha: pre, 9.13 ± 0.55; stim, 11.86 ± 1.11;<br>Beta: pre, 18.3 ± 1; stim, 25.5 ± 2.07; | n = 10 | Two-way ANOVA, F(3, 36) = 12.57, P < 0.0001; Šidák's multiple comparison test. | Pre vs Stim: Delta, p = 0.0006;<br>Theta, p = 0.6823;<br>Alpha, p = 0.4448;<br>Beta: p = 0.0011. |
| Fig.2H | mCherry: no arousal; ChR2: 8.4 ± 1.57 s | n = 10 | Two-tailed unpaired t-test, t = 32.8, df = 18. | p < 0.0001 |
| Fig 2I | mCherry control: no laser: 479.72 ± 27.53 s, 20 Hz: 483.73 ± 31.82 s<br>ChR2 mice: no laser: 512.76 ± 34.2520, 20 Hz: 250.99 ± 16.04 s | mCherry, n = 6; ChR2, n = 7 | Two-way ANOVA, F(1, 11) = 27.73, p = 0.0003; uncorrected Fisher's LSD. | no laser vs 20Hz: mCherry, p = 0.9156; ChR2, p < 0.0001;<br>20Hz: mCherry vs ChR2: p < 0.0001. |
| Fig.2J | Baseline: -9347 ± 4970,<br>2 vol% isoflurane: -517550 ± 17423,<br>2 vol% isoflurane off: -133861 ± 9418 | n = 6 | RM one-way ANOVA, F(1.515, 7.573) = 797.2, p < 0.0001; Šidák's multiple comparison test. | baseline vs 2 vol%, p < 0.0001; 2 vol% vs 2 vol% off, p < 0.0001. |

| | | | | |
|---|---|---|---|---|
| Fig.2L | -30 - 0: no laser, 0.3 ± 1.33; laser, 3.08 ± 0.74;<br>0 - 30: no laser, 7046.22 ± 2556.14; laser, 23141.96 ± 2764.84;<br>30 - 60: no laser, 13003.88 ± 3010.3; laser, 20684.78 ± 14442.65; | n = 6; | Two-way ANOVA, F(1.082, 10.82) = 0.9776, p = 0.352; Tukey's multiple comparison test. | 0 - 30 s: no laser vs laser, p = 0.0016;<br>laser group: -30 - 0 vs 0 - 30, p = 0.0009;<br>no laser group: -30 - 0 vs 30 - 60, p = 0.0172; 0 - 30 vs 30 - 60, p = 0.0052. |
| Fig.2M | no laser, no arousal; laser, 7.17 ± 1.014 s | n = 6; | Two-tailed paired t test, t = 52.11, df = 5. | p < 0.0001 |
| Fig.2N | -420 - -300: no laser, -2.91 ± 1.55; laser, -4.11 ± 1.99;<br>-300 - 0: no laser, 2468.63 ± 39496.08; laser, 203574.83 ± 49686.3;<br>0 - 120: no laser, -114676.02 ± 56048.72; laser, 149239.46 ± 37420.91; | n = 6 | Two-way ANOVA, F(1.462, 14.63) = 6.191, p = 0.0169; Tukey's multiple comparison test. | -300 - 0 s: no laser vs laser, p = 0.0106;<br>0 - 120 s: no laser vs laser, p = 0.0038;<br>laser: -420 - -300 vs -300 - 0, p = 0.0213; -420 - -300 vs 0 - 120, p = 0.0236. |
| Fig.2O | No laser: 97.67 ± 4.65;<br>laser: 112.3 ± 2.704 | n = 6 | Two-tailed paired t test, t = 4.348, df = 5. | p = 0.0074 |
| Fig.2P | -600 - 0: no laser, -6 ± 1.87; laser, -3.14 ± 2;<br>0 - 600: no laser, 701284.62 ± 237169.39; laser, 1699706.5 ± 174589.37; | n = 6 | Two-way ANOVA, F(1, 10) = 11.49, p = 0.0069; uncorrected Fisher's LSD. | 0 - 600 s: no laser vs laser, p = 0.0001;<br>laser: -600 - 0 vs 0 - 600, p < 0.0001;<br>no laser: -600 - 0 vs 0 - 600, p = 0.0071. |
| Fig.2Q | No laser: 465 ± 37.3,<br>laser: 319 ± 38.13 | n = 6; | Two-tailed paired t test, t = 4.052, df = 5. | p = 0.0098 |

| | | | | |
|---|---|---|---|---|
| Fig.3D<br>Total distance of OFT | mCherry:<br>laser off, 3436.75 ± 235.5; laser on, 2929.55 ± 711.95;<br>ChrimsonR: laser off, 4033.67 ± 500.82; laser on, 1434.66 ± 303.01 | n = 6 for mCherry group, n = 7 for ChrimsonR group | Two-way ANOVA, F(1, 11) = 5.047, p = 0.0462; uncorrected Fisher's LSD. | Laser off: mCherry vs ChrimsonR: p = 0.3778;<br>Laser on: mCherry vs ChrimsonR: p = 0.0345;<br>mCherry: laser off vs laser on, p = 0.4734;<br>ChrimsonR: laser off vs laser on, p = 0.0017. |
| Fig.3D<br>The percentage of the time in the center of OFT | mCherry: laser off, 2.375 ± 0.65; laser on, 1.68 ± 0.66;<br>ChrimsonR: laser off, 1.74 ± 0.44; laser on, 0.06 ± 0.06 | n = 6 for mCherry group, n = 7 for ChrimsonR group | Two-way ANOVA, F(1, 11) = 4.198, p = 0.0651; uncorrected Fisher's LSD. | Laser off: mCherry vs ChrimsonR: p = 0.3636;<br>Laser on: mCherry vs ChrimsonR: p = 0.03;<br>mCherry: laser off vs laser on, p = 0.08;<br>ChrimsonR: laser off vs laser on, p = 0.0003. |
| Fig.3D<br>Total Freezing% of the OFT | mCherry: laser off, 13.91 ± 4.08; laser on, 9.27 ± 7.78;<br>ChrimsonR: laser off, 6.77 ± 1.67;<br>laser on, 16.45 ± 7.16 | n = 6 for mCherry group, n = 7 for ChrimsonR group | Two-way ANOVA, F(1, 11) = 2.018, p = 0.1831; uncorrected Fisher's LSD. | Laser off: mCherry vs ChrimsonR: p = 0.3836;<br>Laser on: mCherry vs ChrimsonR: p = 0.3809;<br>mCherry: laser off vs laser on, p = 0.543;<br>ChrimsonR: laser off vs laser on, p = 0.1853. |
| Fig.3D<br>Corner Entropy of the OFT | mCherry: laser off, 1.31 ± 0.25; laser on, 1.22 ± 0.32;<br>ChrimsonR: laser off, 1.3 ± 0.12; | n = 6 for mCherry group, n = 7 for | Two-way ANOVA, F(1, 11) = 13.15, p = 0.004; uncorrected Fisher's LSD. | Laser off: mCherry vs ChrimsonR: p = 0.975; |

| | | | | |
|---|---|---|---|---|
| | laser on, 0.28 ± 0.12 | ChrimsonR group | | Laser on: mCherry vs ChrimsonR: $p = 0.004$;<br>mCherry: laser off vs laser on, $p = 0.64$;<br>ChrimsonR: laser off vs laser on, $p = 0.0001$. |
| Fig.3G<br>Time in laser-on chamber | mCherry: 51.48 ± 2.57;<br>ChrimsonR: 84.57 ± 5.81 | n = 6 for mCherry group, n = 7 for ChrimsonR group | Two-tailed unpaired t test with Welch's correction, $t = 5.21$, $df = 8.205$. | $p = 0.0007$ |
| Fig.3G<br>Distance in laser-on chamber | mCherry: 48.94 ± 1.01;<br>ChrimsonR: 73.66 ± 7.85 | n = 6 for mCherry group, n = 7 for ChrimsonR group | Two-tailed unpaired t test with Welch's correction, $t = 3.122$, $df = 6.2$. | $p = 0.0197$ |
| Fig.3J | Baseline: 44.32 ± 4.72;<br>Test: 41.54 ± 2.55 | n = 7 | Two-tailed paired t test, $t = 0.5883$, $df = 6$. | $p = 0.5778$ |
| Fig.4B | $Ox1R^+$ neurons, 23.35 ± 1.48%; $Ox2R^+$ neurons, 55 ± 3.486%; $vGAT^+$ neurons, 53.39 ± 4.08% | n = 8 slices from 4 brains | Ordinary one-way ANOVA $F(2, 21) = 30.77$, $p < 0.0001$; Šídák's multiple-comparisons test. | Ox1R vs Ox2R, $p < 0.0001$; Ox1R vs vGAT, $p < 0.0001$; Ox2R vs vGAT, $p = 0.9795$. |
| Fig.4D | -30 - 0: no laser, -2.95 ± 2; laser, -4.17 ± 2.45;<br>0 - 30: no laser, -2249.38 ± 543.21; laser, 90094.12 ± 21520.64;<br>30 - 60: no laser, -2431.43 ± 1800.77; laser, 62579.73 ± 17045.44; | n = 7; | Two-way ANOVA, $F(1.476, 17.72) = 14.86$, $p = 0.0004$; Tukey's multiple comparison test. | 0 - 30 s: no laser vs laser, $p = 0.0051$;<br>30 - 60 s: no laser vs laser, $p = 0.0087$;<br>No laser group: -30 - 0 vs 0 - 30, $p = 0.0143$; 0 - 30 vs 30 - 60, $p = 0.9955$;<br>Laser group: -30 - 0 vs 0 - 30, $p = 0.0136$; -30 - 0 vs |

| | | | | |
|---|---|---|---|---|
| | | | | 30 - 60, p = 0.0243; 0 - 30 vs 30 - 60, p = 0.1375. |
| Fig.4E | no laser, no arousal; laser, 8.29 ± 1.73 s | n = 7; | Two-tailed paired t test; t = 29.93, df = 6. | p < 0.0001 |
| Fig.4F | -5 - 0: no laser, 1.83 ± 1.88; laser, -2.3 ± 2;<br>0 - 5: no laser, -111773.88 ± 28183.88; laser, 440355.74 ± 69035.61;<br>5 - 10 (mins): no laser, -176537.55 ± 54809.82; laser, 280504.98 ± 131891.74; | n = 7 | Two-way ANOVA, F(1.409, 16.91) = 13.14, p = 0.0009; Tukey's multiple comparisons test. | laser group:<br>-300 - 0 vs 0 - 300, p = 0.0017;<br>0 - 300 s: no laser vs laser, p < 0.0001, 300 - 600 s: no laser vs laser, p = 0.0126. |
| Fig.4G | -420 - -300: no laser, 3.61 ± 1.5; laser, -2.66 ± 4.24;<br>-300 - 0: no laser, -20250.44 ± 13619.89; laser, 70810.93 ± 25572.71;<br>0 - 120: no laser, 3900.63 ± 7662.14; laser, 29582.38 ± 10117.22; | n = 7 | Two-way ANOVA, F(1.251, 15.01) = 8.704, p = 0.0071; Tukey's multiple comparison test. | -300 - 0 s: no laser vs laser, p = 0.0116;<br>0 - 120 s: no laser vs laser, p = 0.0676;<br>laser: -420 - -300 vs -300 - 0, p = 0.0726; -420 - -300 vs 0 - 120, p = 0.0598. |
| Fig.4H | No laser, 89.71 ± 5.135;<br>laser, 124.7 ± 7.791 s | n = 7 | Two-tailed paired t test; t = 8.08, df = 6. | p = 0.0002 |
| Fig.4I | -600 - 0: no laser, 0.442 ± 1.94; laser, -3.91 ± 4.12;<br>0 - 600: no laser, 1407207.66 ± 654216.92; laser, 2921343.19 ± 791555.64; | n = 7 | Two-way ANOVA, F(1, 12) = 2.174, p = 0.1661; uncorrected Fisher's LSD. | 0 - 600 s: no laser vs laser: p = 0.0479;<br>Laser: -600 - 0 vs 0 - 600 s, p = 0.0017;<br>no laser: -600 - 0 vs 0 - 600, p = 0.0765. |

| | | | | |
|---|---|---|---|---|
| Fig.4J | No laser, 507.6 ± 27.54; laser, 291.6 ± 33.35 s | n = 7 | Two-tailed paired t test; t = 5.172, df = 6. | p = 0.0021 |
| Fig.5B | -30 - 0: no laser, 0.662 ± 3.36; laser, -0.56 ± 2.75; 0 - 30: no laser, -3257.58 ± 1515.18; laser, 25556 ± 4928.3; 30 - 60: no laser, -4930.75 ± 2924.24; laser, 23761.51 ± 16216.96 | n = 6; | Two-way ANOVA, F(1.092, 10.92) = 2.499, p = 0.1414; Tukey's multiple comparisons test. | 0 - 30 s: no laser vs laser, p = 0.0014; 30 - 60 s: no laser vs laser, p = 0.1385; No laser group: -30 - 0 vs 0 - 30, p = 0.1733; 0 - 30 vs 30 - 60, p = 0.677; Laser group: -30 - 0 vs 0 - 30, 0.0081; 0 - 30 vs 30 - 60, p = 0.9951. |
| Fig.5C | no laser: no arousal; laser group: 18 ± 3.95 s | n = 6; | Two-tailed paired t test; t = 10.63, df = 5. | p = 0.0001 |
| Fig.5D | -5 - 0: no laser, 1.81 ± 2.12; laser, 1.78 ± 3.11; 0 - 5: no laser, -38833.62 ± 29746.47; laser, 75169.91 ± 48575.03; 5 - 10 (mins): no laser, -61705.08 ± 64210.69; laser, 76669.44 ± 63950.68 | n = 6; | Two-way ANOVA, F(1.618, 16.18) = 1.808, p = 0.1983; Tukey's multiple comparisons test. | laser group: -5 - 0 vs 0 - 5 mins, p = 0.3478; 0 - 5 mins: no laser vs laser, p = 0.0791; 5 - 10 mins: no laser vs laser, p = 0.1578. |
| Fig.5E | -420 - -300: no laser, 0.563 ± 1.57; laser, -2.42 ± 0.67; -300 - 0: no laser, -32555.66 ± 6305.76; laser, 69330.17 ± 39770.21; 0 - 120: no laser, 1293.65 ± 6254.84; | n = 6; | Two-way ANOVA, F(1.151, 11.51) = 6.145, p = 0.0265; Tukey's multiple-comparisons test. | -300 - 0 s: no laser vs laser, p = 0.0502; 0 - 120 s: no laser vs laser, p = 0.2052; laser: -420 - -300 vs -300 - 0, p = 0.2788; -420 - -300 |

| | | | | |
|---|---|---|---|---|
| | laser, 35546.35 ± 23138.05 | | | vs 0 - 120, p = 0.3521. |
| Fig.5F | No laser, 83 ± 7.759;<br>laser, 120.7 ± 12.55 s | n = 6 | Two-tailed paired t test; t = 3.405, df = 5. | p = 0.0191 |
| Fig.5G | -600 - 0: no laser, 1.14 ± 2.32; laser, 2.24 ± 1.2;<br>0 - 600: no laser, 1686551.32 ± 472635.21; laser, 2825984.67 ± 447374.54 | n = 6; | Two-way ANOVA, F(1, 10) = 3.065, p = 0.1105; uncorrected Fisher's LSD. | 0 - 600 s: no laser vs laser, p = 0.0223;<br>Laser: -600 - 0 vs 0 - 600 s, p = 0.0001;<br>no laser: -600 - 0 vs 0 - 600, p = 0.0044. |
| Fig.5H | No laser, 427.3 ± 13.14;<br>laser, 321.8 ± 23 s | n = 6; | Two-tailed paired t test; t = 3.852, df = 5. | p = 0.012 |
| Fig.5K<br>Total distance | laser off, 5570 ± 1138;<br>laser on, 4012 ± 805.9; | n = 6; | Two-tailed paired t test, t = 3.179, df = 5. | p = 0.0246 |
| Fig.5K<br>the time in the center | laser off, 3.72 ± 1.06;<br>laser on, 0.72 ± 0.17; | n = 6; | Two-tailed paired t test, t = 3.044, df = 5. | p = 0.0286 |
| Fig.5K<br>Total Freezing% | laser off, 23.84 ± 10.36;<br>laser on, 28.65 ± 6.76; | n = 6; | Two-tailed paired t test, t = 0.9224, df = 5. | p = 0.3987 |
| Fig.5K<br>Corner Entropy | laser off, 1.83 ± 0.06;<br>laser on, 1.69 ± 0.08; | n = 6; | Two-tailed paired t test, t = 1.564, df = 5. | p = 0.1785 |
| Fig.5M<br>Time in laser-on chamber | laser off, 55.8 ± 1.79;<br>laser on, 84.3 ± 3.4; | n = 6 | Two-tailed unpaired t test with Welch's correction, t = 7.432, df = 7.567. | p < 0.0001 |
| Fig.5M | mCherry: 52.33 ± 1.91; | n = 6 | Two-tailed unpaired t test with Welch's correction, | p = 0.0003 |

| | | | | |
|---|---|---|---|---|
| Distance in laser-on chamber | ChrimsonR: 78.03 ± 3.68 | | t = 6.208, df = 7.505. | |
| Fig.5O<br>Time in laser-on chamber | Baseline: 42.13 ± 2.63;<br>Test: 40.23 ± 3.09 | n = 6 | Two-tailed paired t test, t = 0.686, df = 5. | p = 0.5232 |
| Fig.6B | -30 - 0: no laser, -3.87 ± 1.63; laser, -3.67 ± 1.68;<br>0 - 30: no laser, 3685.04 ± 1021.94; laser, 25022.25 ± 4829.14;<br>30 - 60: no laser, 952.27 ± 3469.38; laser, 7465.66 ± 2468.79 | n = 6; | Two-way ANOVA, F(1.596, 15.96) = 8.517, p = 0.0047; Tukey's multiple comparisons test. | 0-30 s: no laser vs laser, p = 0.0062,<br>30 - 60 s: no laser vs laser, p = 0.1603;<br>No laser group: -30 - 0 vs 0 - 30, p = 0.0346; 0 - 30 vs 30 - 60, p = 0.7201;<br>laser: -30 - 0 vs 0 - 30 s, p = 0.0081; 0 - 30 vs 30 - 60, p = 0.0528. |
| Fig.6C | no laser: no arousal;<br>laser group: 18.5 ± 1.96 s; | n = 6; | Two-tailed paired t test, t = 21.15, df = 5. | p < 0.0001 |
| Fig.6D | -5 - 0: no laser, 0.29 ± 2.72; laser, 0.44 ± 2.81;<br>0 - 5: no laser, 7652.11 ± 27289.22; laser, 149308.2 ± 52117.65;<br>5 - 10 (mins): no laser, -42205.68 ± 56290.62; laser, 144390.75 ± 79043.75 | n = 6; | Two-way ANOVA, F(1.490, 14.90) = 3.143, p = 0.0839; Tukey's multiple comparisons test. | laser group: -5 - 0 vs 0 - 5 mins, p = 0.0765;<br>0 - 5 mins: no laser vs laser, p = 0.0444,<br>5 - 10 mins: no laser vs laser, p = 0.0865. |
| Fig.6E | -420 - -300: no laser, 5.39 ± 1.26; laser, -1.38 ± 3.18;<br>-300 - 0: no laser, -1448.55 ± 31933.63; laser, -7344.17 ± 19894.26; | n = 6; | Two-way ANOVA, F(1.292, 12.92) = 0.04263, p = 0.8939; Tukey's multiple comparisons test. | -300 - 0 s: no laser vs laser, p = 0.8792;<br>0 - 120 s: no laser vs laser, p = 0.7051;<br>laser: -420 - -300 vs -300 - 0, p = |

| | | | | |
|---|---|---|---|---|
| | 0 - 120: no laser, 25982.77 ± 16087.28; laser, 17900.26 ± 13073.33 | | | 0.9287; -420 - -300 vs 0 - 120, p = 0.4223. |
| Fig.6F | no laser, 95.17 ± 9.58,<br>laser, 117 ± 12.17 s; | n = 6; | Two-tailed paired t test, t = 1.763, df = 5. | p = 0.1382 |
| Fig.6G | -600 - 0: no laser, 1.65 ± 3.75; laser, 4.29 ± 4.06<br>0 - 600: no laser, 1649355.05 ± 351579.31; laser, 1781995.77 ± 632144.63 | n = 6; | Two-way ANOVA, F(1, 10) = 0.03362, p = 0.8582; uncorrected Fisher's LSD. | 0 - 600 s: no laser vs laser, p = 0.798;<br>Laser: -600 - 0 vs 0 - 600 s, p = 0.0059;<br>no laser: -600 - 0 vs 0 - 600, p = 0.0091. |
| Fig.6H | no laser: 512.8 ± 39.38,<br>laser: 433.2 ± 17.48 s; | n = 6; | Two-tailed paired t test, t = 3.001, df = 5. | p = 0.0301 |
| Fig.6K<br>Total distance | laser off, 6067 ± 1060;<br>laser on, 6621 ± 1784; | n = 8 | Two-tailed paired t test, t = 0.6305, df = 7. | p = 0.5484 |
| Fig.6K<br>the time in the center | laser off, 3.81 ± 1.147;<br>laser on, 2.07 ± 0.68; | n = 8 | Two-tailed paired t test, t = 2.33, df = 7. | p = 0.0526 |
| Fig.6K<br>Total Freezing% | laser off, 19.39 ± 4.35;<br>laser on, 19.11 ± 4.43; | n = 8 | Two-tailed paired t test, t = 0.0783, df = 7. | p = 0.9398 |
| Fig.6K<br>Corner Entropy | laser off, 1.76 ± 0.04;<br>laser on, 1.36 ± 0.24; | n = 8 | Two-tailed paired t test, t = 1.774, df = 7. | p = 0.1194 |
| Fig.6M<br>Time in laser-on chamber | laser off, 40.5 ± 4.34;<br>laser on, 16.51 ± 5.09; | n = 5 | Two-tailed unpaired t test with Welch's correction, t = 3.586, df = 7.802. | p = 0.0074 |

| | | | | |
|---|---|---|---|---|
| Fig.6M<br>Distance in laser-on chamber | mCherry: 45.86 ± 2.2;<br>ChrimsonR: 23.77 ± 4.14 | n = 5 | Two-tailed unpaired t test with Welch's correction, t = 4.716, df = 6.087. | p = 0.0031 |
| Fig.6O | Baseline: 48.12 ± 5.61;<br>Test: 37.18 ± 3.37 | n = 6 | Two-tailed paired t test, t = 1.734, df = 5. | p = 0.1435 |
| Supplementary Fig.1B<br>IPSCs amplitude with Ox1R antagonist SB-334867 | Before: 60.31 ± 4.19, SB-334867: 49.57 ± 10.03, Washout: 61.4 ± 8.11 | n=10 | Repeated measures one-way ANOVA, F(1.361, 12.25) = 1.384, p = 0.2753, Šídák's multiple comparisons. | baseline vs SB-334867: p = 0.6673, SB-334867 vs washout: p = 0.1215. |
| Supplementary Fig.1B<br>IPSCs amplitude with Ox2R antagonist TCS Ox2 29 | Before: 62.48 ± 4.23, TCS Ox2 29: 38.4 ± 7.22, Washout: 62.13 ± 5.67. | n=10 | Repeated measures one-way ANOVA, F(1.434, 12.91) = 6.065, p = 0.0203, Šídák's multiple comparisons. | baseline vs TCS OX2 29: p = 0.0552, TCS OX2 vs washout: p = 0.1029. |
| Supplementary Fig.2B<br>EEG spectral power in mCherry mice | Delta: pre, 34.89 ± 1.1; stim, 36.71 ± 1.22;<br>Theta: pre, 38.22 ± 1.21; stim, 35.89 ± 1.68;<br>Alpha: pre, 16.19 ± 0.53; stim, 15.82 ± 1.25;<br>Beta: pre, 11.24 ± 0.5; stim, 12 ± 0.41; | n = 7 | Two-way ANOVA, F(3, 24) = 2.533, p = 0.0809, Šídák's multiple comparisons. | Pre vs Stim:<br>Delta, p = 0.3881, Theta, p = 0.1773, Alpha, p = 0.9956, Beta, p = 0.9383. |
| Supplementary Fig.2B<br>EEG spectral power in ChR2 mice | Delta: pre, 35.65 ± 1.93; stim, 21.17 ± 1.63;<br>Theta: pre, 33.56 ± 2; stim, 41.23 ± 2.67;<br>Alpha: pre, 15.97 ± 1.73; stim, 14.04 ± 1.22; | n = 7 | Two-way ANOVA, F(3, 24) = 30.1, p < 0.0001, Šídák's multiple comparisons. | Pre vs Stim:<br>Delta, p < 0.0001, Theta, p = 0.0026, Alpha, p = 0.8025, Beta, p = 0.0008; |

| | | | | |
|---|---|---|---|---|
| | Beta: pre, 15.16 ± 1.1; stim, 23.74 ± 1.2 | | | |
| Supplementary Fig.2C | ChR2:<br>5 Hz, 13.57 ± 2.94; 10 Hz, 10.57 ± 2.44; 20 Hz, 5.71 ± 1.55; 30 Hz, 7.14 ± 1.6 | n = 7 | Two-way ANOVA, F(1.566, 18.79) = 14.91, p = 0.0003, Tukey's multiple comparisons. | mCherry vs ChR2:<br>5 Hz, p < 0.0001,<br>10 Hz, p < 0.0001,<br>20 Hz, p < 0.0001,<br>30 Hz, p < 0.0001. |
| Supplementary Fig.2D | mCherry group: no laser: 10.6 ± 0.7 s vs. laser: 11.28 ± 0.9 s;<br>ChR2 group: no laser: 10.61 ± 0.44 s vs. laser 10.56 ± 0.59 s | mCherry group: n = 6; ChR2 group: n = 8 | Two-way ANOVA, F(1,12) = 0.5267, p = 0.4819; uncorrected Fisher's LSD. | mCherry, no laser vs laser: p = 0.3886;<br>ChR2, no laser vs laser: p = 0.941. |
| Supplementary Fig.3B | -30 - 0: no laser, 3.3 ± 2.01; laser, 0.422 ± 0.88;<br>0 - 30: no laser, 4308.91 ± 2429.57; laser, -234.87 ± 1374.56;<br>30 - 60: no laser, 2080.94 ± 3141.73; laser, -1466.76 ± 904.76; | n = 5 | Two-way ANOVA, F(1.924, 15.4) = 1.171, p = 0.3344; Tukey's multiple comparison test. | No laser group:<br>-30 - 0 vs 0 - 30, p = 0.2891; -30 - 0 vs 30 - 60, p = 0.7967; 0 - 30 vs 30 - 60, p = 0.7768;<br>laser group:<br>-30 - 0 vs 0 - 30, p = 0.984; -30 - 0 vs 30 - 60, p = 0.3376; 0 - 30 vs 30 - 60, p = 0.3194. |
| Supplementary Fig.3C | No arousal behaviors in both no laser and laser conditions | n = 5 | | |
| Supplementary Fig.3D | -420 - -300: no laser, 4.73 ± 3.48; laser, 0.582 ± 8.53;<br>-300 - 0: no laser, 20603.902 ± 31818.69; laser, 77246.27 ± 49741.32;<br>0 - 120: no laser, -45135.76 ± 29798.34; | n = 5 | Two-way ANOVA, F(1.872, 14.97) = 12.28, p = 0.0008; Tukey's multiple comparison test. | -300 - 0 s: no laser vs laser, p = 0.37; 0 - 120 s: no laser vs laser, p = 0.007;<br>no laser: -420 - -300 vs -300 - 0, p = 0.804; -420 - -300 vs 0 - 120, p = 0.3764; -300 - 0 |

| | | | | |
|---|---|---|---|---|
| | laser, -291094.02 ± 54552.47; | | | vs 0 - 120, p = 0.046;<br>laser: -420 - -300 vs -300 - 0, p = 0.362; -420 - -300 vs 0 - 120, p = 0.013; -300 - 0 vs 0 - 120, p = 0.014. |
| Supplementary Fig.3E | No laser: 106.2 ± 2.85; laser: 103 ± 2.76 | n = 5; | Two-tailed paired t test, t = 0.651, df = 4. | p = 0.5504 |
| Supplementary Fig.3F | -600 - 0: no laser, -2.09 ± 4.27; laser, 0.59 ± 2.58;<br>0 - 600: no laser, 1831513.48 ± 366314.13; laser, 1430700.7 ± 419444.53; | n = 5 | Two-way ANOVA, F(1, 8) = 0.518, p = 0.4922; uncorrected Fisher's LSD. | 0 - 600 s: no laser vs laser, p = 0.3239;<br>laser: -600 - 0 vs 0 - 600, p = 0.0067;<br>no laser: -600 - 0 vs 0 - 600, p = 0.0016. |
| Supplementary Fig.3G | No laser, 357 ± 13.83;<br>Laser, 400.2 ± 27.33 | n = 5 | Two-tailed paired t test, t = 1.691, df = 4. | p = 0.1661 |
| Supplementary Fig.4B | $Ox1R^+$ neurons, 18.4 ± 1.16%; $Ox2R^+$ neurons, 50.06 ± 4.37%; $TH^+$ neurons, 5.58 ± 0.85% | n = 8 slices from 4 brains | Ordinary one-way ANOVA, F(2, 21) = 74.23, p < 0.0001; Šídák's multiple-comparisons test. | Ox1R vs Ox2R, p < 0.0001; Ox1R vs TH, p = 0.0079; Ox2R vs TH, p < 0.0001. |
| Supplementary Fig.4D | $Ox1R^+$ neurons, 18.3 ± 1.79%; $Ox2R^+$ neurons, 57.51 ± 3.64%; $vGlut2^+$ neurons, 24.81 ± 3.18% | n = 8 slices from 4 brains | Ordinary one-way ANOVA, F(2, 21) = 49.83, p < 0.0001; Šídák's multiple-comparisons test. | Ox1R vs Ox2R, p < 0.0001; Ox1R vs vGlut2, p = 0.356; Ox2R vs vGlut2, p < 0.0001. |
| Supplementary Fig.4F | -30 - 0: no laser, 1.94 ± 1.3; laser, -0.91 ± 1.88;<br>0 - 30: no laser, 2438.99 ± 1059.47; laser, 1265.7 ± 450.41;<br>30 - 60: no laser, 1854.21 ± 1293.2; | n = 10; | Two-way ANOVA, F(1.458, 26.24) = 0.5363, p = 0.536; Tukey's multiple comparisons test. | 0 - 30 s: no laser vs laser, p = 0.328;<br>30 - 60 s: no laser vs laser, p = 0.7587;<br>No laser group: -30 - 0 vs 0 - 30, p = |

| | | | | |
|---|---|---|---|---|
| | laser, 2525.59 ± 1717.36; | | | 0.1071; 0 - 30 vs 30 - 60, p = 0.7795;<br>Laser group: -30 - 0 vs 0 - 30, p = 0.0482; 0 - 30 vs 30 - 60, p = 0.7584. |
| Supplementary Fig.4G | no laser, no arousal;<br>laser, 10.6 ± 1.57 s | n = 10; | Two-tailed paired t test; t = 31.43, df = 9. | p < 0.0001 |
| Supplementary Fig.5B | -30 - 0: no laser, 3.23 ± 1.31; laser, -0.84 ± 1.59;<br>0 - 30: no laser, -795.75 ± 1645.74; laser, -1766.51 ± 1319.12;<br>30 - 60: no laser, -3848.53 ± 3171.57; laser, 80.07 ± 3172.66; | n = 6; | Two-way ANOVA, F(1.346, 13.46) = 1.014, p = 0.3577; Tukey's multiple-comparisons test. | 0 - 30 s: no laser vs laser, p = 0.6556;<br>30 - 60 s: no laser vs laser, p = 0.4017;<br>No laser group: -30 - 0 vs 0 - 30, p = 0.8812; 0 - 30 vs 30 - 60, p = 0.4924;<br>Laser group: -30 - 0 vs 0 - 30, p = 0.4366; 0 - 30 vs 30 - 60, p = 0.8189. |
| Supplementary Fig.5C | no arousal in both no laser and laser group | n = 6; | | |
| Supplementary Fig.5D | -5 - 0: no laser, 1.41 ± 3.03; laser, -0.11 ± 1.77;<br>0 - 5: no laser, -74768.06 ± 52554.34; laser, -20700.14 ± 31226.84;<br>5 - 10 (mins): no laser, -114665.55 ± 93007.68; laser, -111400.87 ± 69642.28; | n = 6 | Two-way ANOVA with Tukey's multiple-comparisons test, F(1.059, 10.59) = 0.2645, p = 0.6311. | laser group: -300 - 0 vs 0 - 300, p = 0.7938;<br>0 - 300 s: no laser vs laser, p = 0.4018,<br>300 - 600 s: no laser vs laser, p = 0.9782. |
| Supplementary Fig.5E | -420 - -300: no laser, -4.04 ± 3.42; laser, 1.21 ± 2.65; -300 - 0: no laser, 5355.15 ± 15479.12; laser, | n = 6 | Two-way ANOVA with Tukey's multiple-comparisons test, | -300 - 0 s: no laser vs laser, p = 0.6254; |

| | | | | |
|---|---|---|---|---|
| | 23314.09 ± 31656.93; 0 - 120: no laser, -147.6 ± 10303.02; laser, 9970.54 ± 13228.89 | | F(1.346, 13.46) = 0.2011, p = 0.7328. | 0 - 120 s: no laser vs laser, p = 0.5604;<br>laser: -420 - -300 vs -300 - 0, p = 0.7543; -420 - -300 vs 0 - 120, p = 0.7449. |
| Supplementary Fig.5F | No laser, 137 ± 13.83;<br>laser, 117.5 ± 11.95 s | n = 6 | Two-tailed paired t test; t = 2.052, df = 5. | p = 0.0954 |
| Supplementary Fig.5G | -600 - 0: no laser, 2.51 ± 1.99; laser, 1.17 ± 1.23;<br>0 - 600: no laser, 778229.73 ± 304059; laser, 794396.18 ± 131346.08 | n = 6; | Two-way ANOVA with uncorrected Fisher's LSD; F(1, 10) = 0.00238, p = 0.962. | 0 - 600 s: no laser vs laser: p = 0.9457;<br>Laser: -600 - 0 vs 0 - 600 s, p = 0.0069;<br>no laser: -600 - 0 vs 0 - 600, p = 0.0077. |
| Supplementary Fig.5H | No laser, 472 ± 75.22;<br>laser, 433.3 ± 86.19 s | n = 6 | Two-tailed paired t test; t = 0.6611, df = 5. | p = 0.5378 |

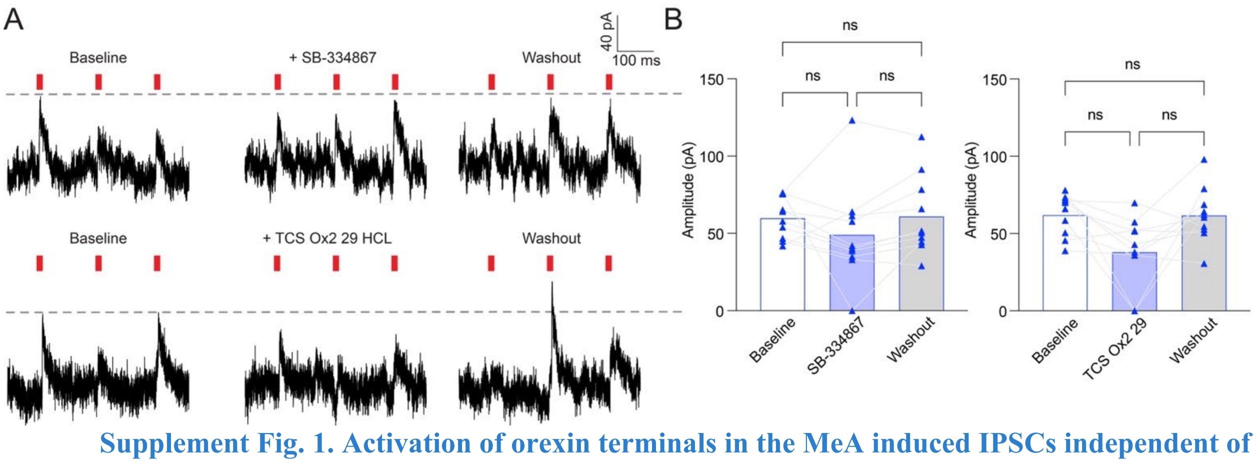


**Supplement Fig. 1. Activation of orexin terminals in the MeA induced IPSCs independent of Ox1R and Ox2R signaling.**

(**A**) Representative traces showed that IPSCs were not blocked by the Ox1R antagonist SB-334867 or the Ox2R antagonist TCS Ox2 29. (**B**) Quantification of IPSC amplitudes before, during, and after blockage. Repeated measures one-way ANOVA with Šídák's multiple-comparisons test, n = 10. Data are mean ± S.E.M.; statistical details and exact n values are provided in Supplementary Table 1.

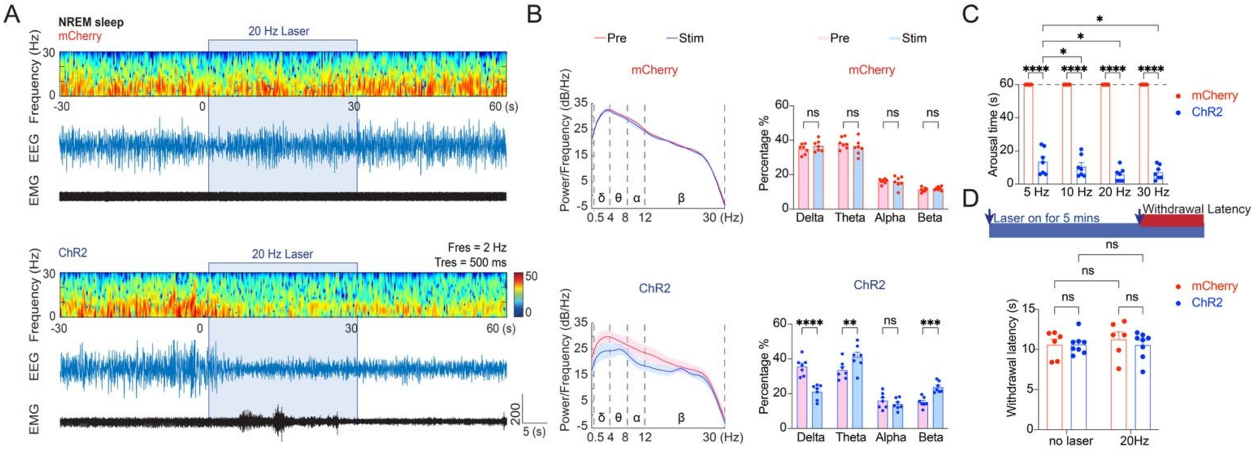


**Supplement Fig. 2. Selective activation of LHA$^{Ox}$→MeA projection promoted wakefulness from NREM sleep but did not induce analgesia.**

(**A**) EEG spectrograms and representative EEG/EMG traces show that 20Hz optogenetic stimulation during NREM sleep induced wakefulness in ChR2 mice but not in mCherry controls. (**B**) EEG power spectral density (PSD) plots and frequency band analysis before and during MeA orexin terminal stimulation. Two-way ANOVA followed by Šídák's multiple-comparisons test (n = 7 for each group). (**C**) Arousal behavioral responses showed that 5Hz, 10Hz, 20Hz, and 30Hz selective optogenetic activation of LHA$^{Ox}$→MeA projection enhanced arousal from NREM sleep in ChR2 but not mCherry mice. Two-way ANOVA, Tukey's multiple comparisons test, n = 7 for each group. (**D**) Schematic diagram of the hotplate test (above). Optogenetic stimulation of LHA$^{Ox}$→MeA projection did not affect the withdrawal latency to 55°C hotplate. Two-way ANOVA, uncorrected Fisher's LSD, n = 6 for mCherry group, n = 8 for ChR2 group. Data are mean ± S.E.M.; statistical details and exact n values are provided in Supplementary Table 1. *$p \leq 0.05$, **$p \leq 0.01$, ***$p < 0.001$, ****$p < 0.0001$.

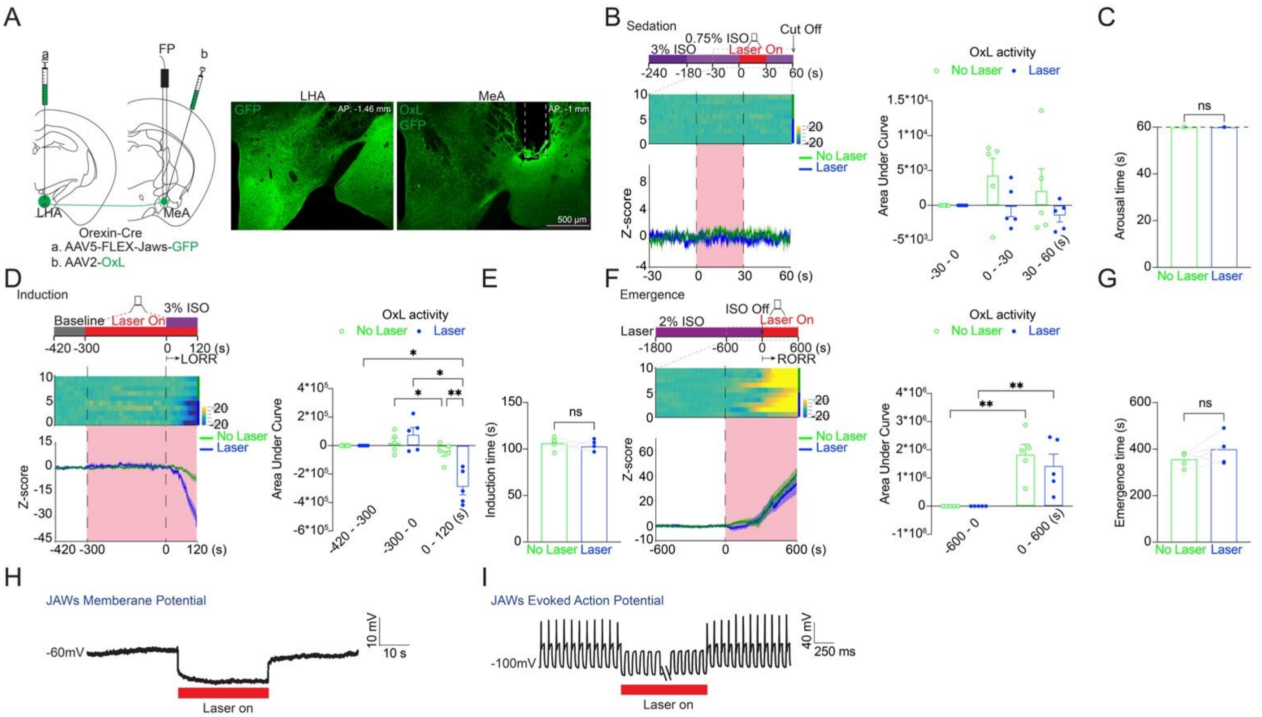


**Supplement Fig. 3. Inhibition of the $\text{LHA}^{\text{Ox}}$→ MeA circuit did not affect Ox release in the MeA during anesthesia induction, duration, and emergence.**

(**A**) Left: Cre-dependent JAWs was injected into the LHA, and orexin sensor Oxlight (OxL) was injected into the MeA, followed by the fiber implantation into the same place. Right: Representative images showed the JAWs and OxL expression. (**B**) Left: Fluorescence trials and Z-score of averaged Oxlight photometry signals from the JAWs group in response to arousal behaviors. Right: The Ox release in the MeA was not altered while the optogenetic inhibition of $\text{LHA}^{\text{Ox}}$→MeA projection during the light anesthesia in Jaws no laser and Jaws laser. Two-way ANOVA, Tukey's multiple comparisons test, n = 5. (**C**) Inhibition of $\text{LHA}^{\text{Ox}}$→MeA projections did not produce arousal from light anesthesia. n = 5. (**D**) Left: Fluorescence trials and Z-score of averaged Oxlight photometry signals showed that pre-inhibition of the $\text{LHA}^{\text{Ox}}$→MeA projection during induction did not affect Ox release. Right: The Z-score of the area under the curve data showed Ox release did not change while inhibiting $\text{LHA}^{\text{Ox}}$→MeA projection before and during anesthesia induction. Two-way ANOVA, Tukey's multiple comparisons test, n = **5**. (**E**) Pre-inhibiting $\text{LHA}^{\text{Ox}}$→MeA projection did not affect the induction time. Two-tailed paired t test for the same group comparison, n = 5. (**F**) Left: Inhibiting the $\text{LHA}^{\text{Ox}}$→MeA projection did not affect the Ox release during anesthesia emergence. Right: The Z-score of the area under the curve data showed Ox release while inhibiting $\text{LHA}^{\text{Ox}}$→MeA projection during emergence from anesthesia. Two-way ANOVA, Uncorrected Fisher's LSD, n = 5 for each group. (**G**) The return of the righting reflex data showed that the inhibition of $\text{LHA}^{\text{Ox}}$→MeA projection did not affect the emergence from anesthesia. Two-tailed paired t test for the same group comparison, n = 5 for each group. (**H** to **I**). Patch-clamp whole-cell recordings on $\text{LHA}^{\text{Ox}}$ neurons showed that optogenetic activation of JAWs decreased the membrane potential (**H**) and silenced the action potentials evoked by injecting 50 pA current at 10 Hz (**I**). Data are mean ± S.E.M.; statistical details and exact n values are provided in Supplementary Table 1. $*p \leq 0.05$, $**p \leq 0.01$.

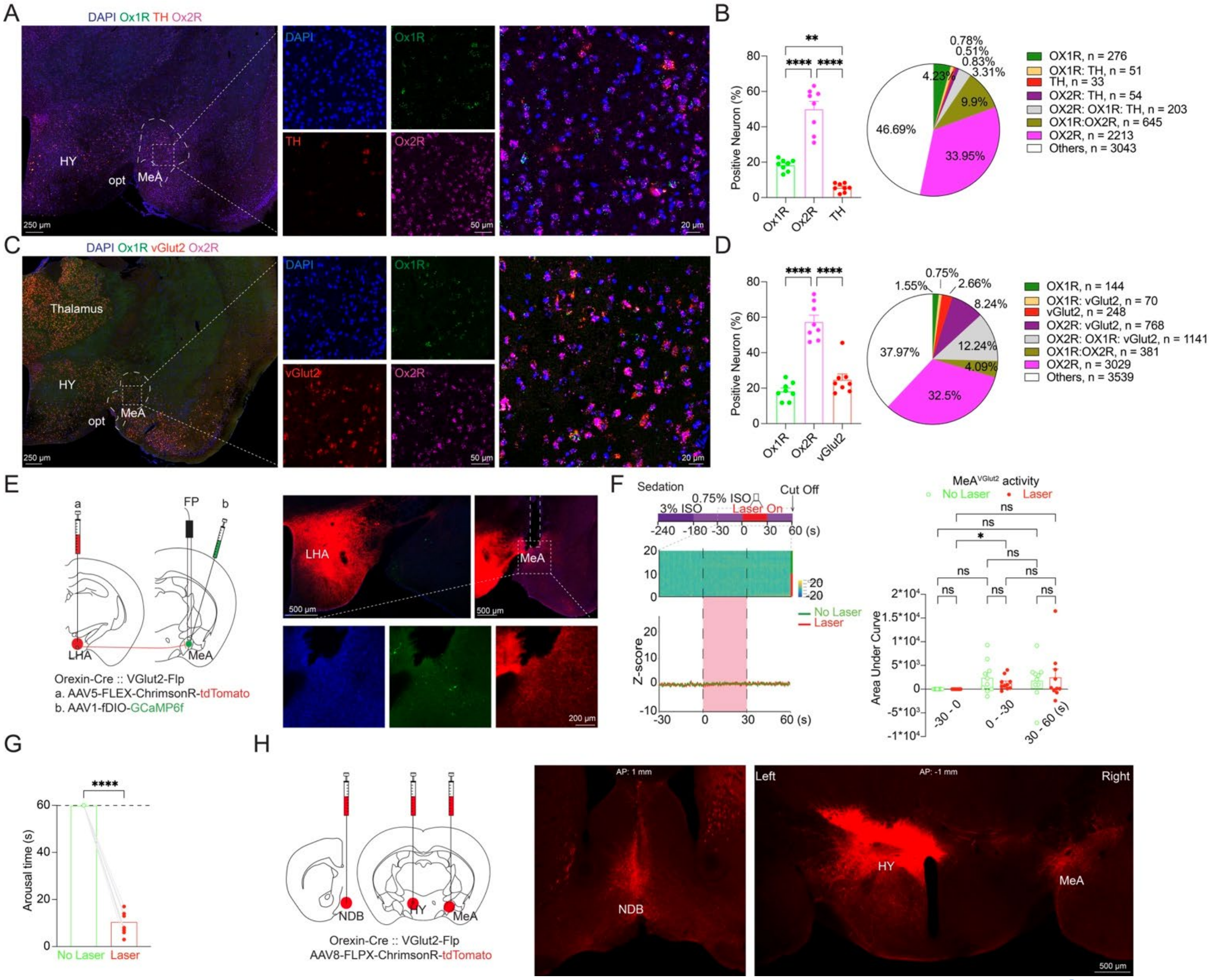


**Supplement Fig. 4. TH and vGlut2 neurons were not major downstream targets of LHA$^{Ox}$→ MeA projection.**

(**A**) Representative RNAscope images showed DAPI (blue), Ox1R (green), TH (red), and Ox2R (magenta) mRNA expression in the MeA. (**B**) Quantification of Ox1R$^{+}$, Ox2R$^{+}$, and TH$^{+}$ cells as a percentage of DAPI-positive cells per slice. Data were analyzed using ordinary one-way ANOVA followed by Šídák's multiple-comparisons test (n = 8 slices from 4 brains). The pie chart summarized the distribution of neuronal populations pooled from all slices. (**C**) Representative RNAscope images showed DAPI (blue), Ox1R (green), vGlut2 (red), and Ox2R (magenta) mRNA expression in the MeA. Scale bars, 250, 50, and 20 µm (bottom right). (**D**) Quantification of Ox1R$^{+}$, Ox2R$^{+}$, and vGlut2$^{+}$ cells as a percentage of DAPI-positive cells per slice. Data were analyzed using ordinary one-way ANOVA followed by Šídák's multiple-comparisons test (n = 8 slices from 4 brains). The pie chart summarized the distribution of neuronal populations pooled from all slices. (**E**) Experimental strategy combining Cre-dependent ChrimsonR expression in the LHA with Flp-dependent GCaMP6f expression in the MeA$^{vGlut2}$ neurons, with an optic fiber implanted above the MeA. Representative images confirmed ChrimsonR expression in the LHA and minimal GCaMP6f expression in the MeA. (**F**) Optogenetic activation of orexin terminals in MeA did not significantly alter the MeA$^{vGlut2}$ neurons' activity during isoflurane sedation. Two-way ANOVA with Tukey's multiple-comparisons test, n = 10. (**G**) Despite the lack of vGlut2 neurons' activation, optogenetic activation of orexin terminals in MeA induced rapid arousal from light anesthesia. n = 10. (**H**) A Flp-dependent virus (AAV8-FLPX-ChrimsonR-tdTomato) was injected in diagonal band nucleus (NDB), MeA, and LHA of the Orexin-Cre::VGlut2-Flp mice. Representative images confirmed robust ChrimsonR expression in the NDB and LHA, but little expression in the MeA. Data are mean ± S.E.M.; statistical details and exact n values are provided in Supplementary Table 1. *$p \leq 0.05$, **$p \leq 0.01$, ****$p < 0.0001$.

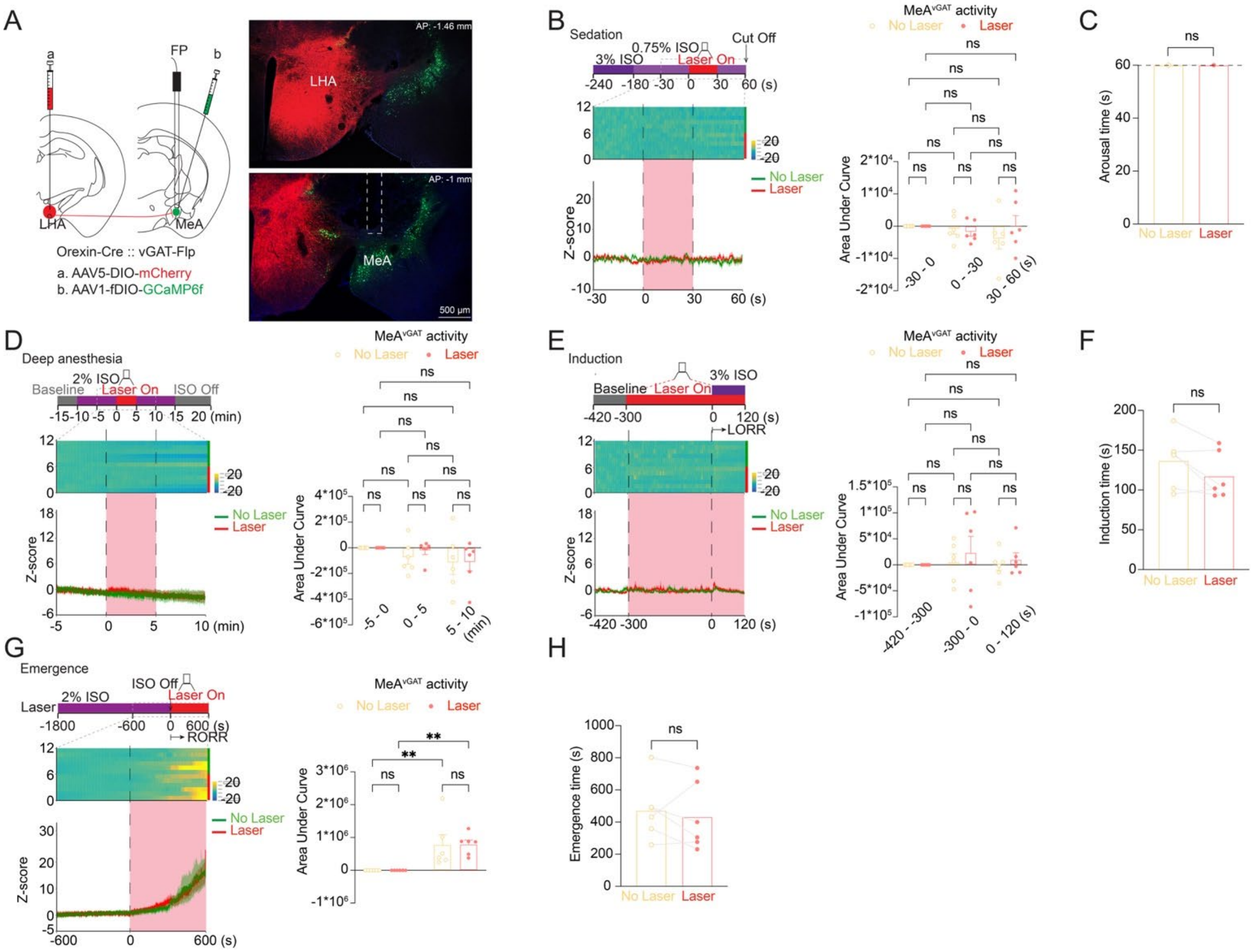


**Supplement Fig. 5. Control experiments showed no effect of orexin terminal activation on MeA$^{vGAT}$ neuron activity or anesthetic behaviors in mCherry mice.**

(**A**) Experimental strategy combining Cre-dependent mCherry expression in LHA$^{Ox}$ neurons with Flp-dependent GCaMP6f expression in MeA$^{vGAT}$ neurons, with an optic fiber implanted above the MeA of the Orexin-Cre::vGAT-Flp mice. Representative images confirmed ChrimsonR and GCaMP6f expression. (**B**, **D**, **E**, and **G**). In the mCherry group, optogenetic activation of orexin terminals in the MeA did not alter MeAvGAT neurons activity during isoflurane sedation (**B**, Two-way ANOVA with Tukey's multiple-comparisons test; n = 6), 2% isoflurane anesthesia (**D**, Two-way ANOVA with Tukey's multiple-comparisons test; n = 6), isoflurane induction (E, Two-way ANOVA with Tukey's multiple-comparisons test; n = 6), or anesthesia emergence (**G**, Two-way ANOVA with Uncorrected Fisher's LSD; n = 6). (**C**, **F**, and **H**). In the mCherry group, optogenetic activation of orexin terminals in the MeA did not induce rapid arousal from light anesthesia (**C**, two-tailed paired t test; n = 6), the time to loss of the righting reflex (LoRR) during anesthesia induction (**F**, two-tailed paired t test; n = 6) and the time to recovery of the righting reflex (RoRR) (**H**, two-tailed paired t test; n = 6). Data are mean ± S.E.M.; statistical details and exact n values are provided in Supplementary Table 1. *$p \leq 0.05$, **$p \leq 0.01$, ****$p < 0.0001$.
